# Nonsense-mediated mRNA decay relies on “two-factor authentication” by SMG5-SMG7

**DOI:** 10.1101/2020.07.07.191437

**Authors:** Volker Boehm, Sabrina Kueckelmann, Jennifer V. Gerbracht, Thiago Britto-Borges, Janine Altmüller, Christoph Dieterich, Niels H. Gehring

**Affiliations:** Institute for Genetics, University of Cologne, 50674 Cologne, Germany; Center for Molecular Medicine Cologne (CMMC), University of Cologne, 50937 Cologne, Germany; Section of Bioinformatics and Systems Cardiology, Department of Internal Medicine III and Klaus Tschira Institute for Integrative Computational Cardiology, Heidelberg University Hospital, 69120 Heidelberg, Germany; DZHK (German Centre for Cardiovascular Research), Partner site Heidelberg/Mannheim, 69120 Heidelberg, Germany; Cologne Center for Genomics (CCG), University of Cologne, 50931 Cologne, Germany

**Keywords:** Nonsense-mediated mRNA decay, mRNA degradation, gene expression, UPF1, SMG5-SMG7, SMG6

## Abstract

Eukaryotic gene expression is constantly regulated and controlled by the translation-coupled nonsense-mediated mRNA decay (NMD) pathway. Aberrant translation termination leads to NMD activation and robust clearance of NMD targets via two seemingly independent and redundant mRNA degradation branches. Here, we uncover that the loss of the first SMG5-SMG7-dependent pathway also inactivates the second SMG6-dependent branch, indicating an unexpected functional hierarchy of the final NMD steps. Transcriptome-wide analyses of SMG5-SMG7-depleted cells confirm complete NMD inhibition resulting in massive transcriptomic alterations. The NMD activity conferred by SMG5-SMG7 is determined to varying degrees by their interaction with the central NMD factor UPF1, heterodimer formation and the initiation of deadenylation. Surprisingly, we find that SMG5 functionally substitutes SMG7 and vice versa. Our data support an improved model for NMD execution that requires two-factor authentication involving UPF1 phosphorylation and SMG5-SMG7 recruitment to access SMG6 activity.

## Introduction

Error-free and precisely regulated gene expression is an essential prerequisite for all living organisms. In eukaryotes, transcription and translation are controlled and fine-tuned by diverse mechanisms to ensure the generation of flawless RNAs and proteins^1^. Mature mRNAs that have completed all co- and post-transcriptional processing steps and passed the associated quality checks are translated into proteins as the final step of gene expression in the cytoplasm. At this point, translation-coupled mechanisms inspect the mRNA one last time to perform a final quality control. Specifically, it is assessed whether the translated mRNAs are legitimate or contain features indicating that these transcripts encode non-functional, incomplete or potentially harmful proteins and therefore have to be degraded^2,3^. The arguably most famous translation-coupled quality control process is nonsense-mediated mRNA decay (NMD), which is best known for its role to remove mutated transcripts containing a premature termination codon (PTC)^4^. However, the relevance of NMD for cellular maintenance goes beyond the quality control function and is not restricted to mutated transcripts^5^. Previous studies found that about 5-10% of the expressed genes are affected by NMD in different organisms^6–15^, suggesting that NMD serves as a regulatory mechanism, which fine-tunes general gene expression and helps to minimize the production of aberrant transcript isoforms. Furthermore, defects in the core NMD machinery are not compatible with life in higher eukaryotes^16–22^, underlining the importance of NMD to function properly during development and cellular maintenance.

In general, inefficient translation termination seems to be the primary stimulus for NMD initiation^4^. Recent evidence suggests that NMD can in principle be triggered by each translation termination event with a certain probability^23^. In higher eukaryotes, this probability can be modulated by different NMD-activating features, such as a long 3’ untranslated region (UTR)^24–27^. However, the exact length and composition of an NMD-activating 3’ UTR is not exactly defined and many mRNAs contain NMD-suppressing sequences that allow them to escape this type of NMD^28–30^. Another potent activator of NMD is the presence of an RNA-binding protein complex called the exon-junction complex (EJC) downstream of a terminating ribosome^24,31–37^. The EJC serves as a mark for successful splicing and is deposited onto the mRNA approximately 20-24 nucleotides upstream of spliced junctions^38–41^. Stop codons are typically located in the last exon of regular protein-coding transcripts, thus ribosomes usually displace all EJCs from a translated mRNA, effectively removing the degradation-inducing feature. However, mutations or alternative splicing may produce isoforms with stop codons situated upstream of EJC deposition sites. Translation of these transcripts would fail to remove all EJCs and subsequently triggers the decay of the mRNA via efficiently activated NMD.

Intensive research over many decades uncovered the central players of the complex NMD pathway and how they cooperate to achieve highly specific and efficient mRNA degradation. According to generally accepted models, NMD execution requires a network of factors to identify a given translation termination event as aberrant^42^. The RNA helicase UPF1 holds a central position in the NMD pathway, as it serves as a binding hub for other NMD factors and is functionally involved in all stages from the recognition of NMD substrates until the disassembly of the NMD machinery^43^. In translation-inhibited conditions, UPF1 has the potential to bind non-specifically to all expressed transcripts. However, in unperturbed cells, UPF1 is found preferentially in the 3′ UTR region of translated mRNA due to the displacement from the 5′ UTR and coding region by translating ribosomes^44–47^. Furthermore, the ATPase and helicase activity of UPF1 is required to achieve target discrimination, resulting in increased binding of NMD-targets and release of UPF1 from non-target mRNAs^48^.

If the translated mRNA contains a premature or otherwise aberrant termination codon, the downstream bound UPF1 promotes the recruitment of NMD factors and their assembly into an NMD-activating complex. Subsequently, protein-protein interactions between UPF1, UPF2, UPF3 and - if present - the EJC stimulate the phosphorylation of SQ/TQ motifs in UPF1 by the kinase SMG1^49–53^. Importantly, the activity of the kinase SMG1 is regulated by multiple accessory NMD factors, presumably to prevent unwanted UPF1 phosphorylation on non-NMD targets^54–58^. Continued presence of UPF1 on the target transcript in an NMD-activating environment leads to gradually increasing hyper-phosphorylation of UPF1 at up to 19 potential phosphorylation sites^59^. The progressively phosphorylated residues in the N- and C-terminal tails of UPF1 then act as binding sites for the decay-inducing factors SMG5, SMG6 and SMG7^60^. In this basic model, hyper-phosphorylation of UPF1 represents a “point of no return” for NMD activation, which effectively sentences the mRNA for degradation^61^.

The final execution of NMD is divided into two major branches. The first branch relies on the interaction of phosphorylated UPF1 with the heterodimer SMG5-SMG7, which in turn recruits the CCR4-NOT deadenylation complex^62–64^. Consequently, SMG5-SMG7 promote target mRNA deadenylation, followed by decapping and 5′-3′ or 3′-5′ exonucleolytic degradation^65,66^. The second branch is mediated by the endonuclease SMG6, which interacts with UPF1 to cleave the NMD-targeted transcript in a region around the NMD-activating stop codon^36,67–71^. This endonucleolytic cleavage results in the generation of two decay intermediates, which are rapidly removed by exonucleolytic decay. Both SMG5-SMG7 and SMG6-mediated degradation pathways are considered to be redundant, as they target the same transcripts^14^. They are also regarded as independent, because downregulation of individual factors (SMG5, SMG6 or SMG7) only partially inhibits NMD^35,62,72^. However, loss of SMG6 impaired NMD more severely than inactivation of SMG7, suggesting that endonucleolytic cleavage is the preferred decay pathway, whereas deadenylation has merely a backup/supplementary function^14,35,62,70,71,73^. Nonetheless, the apparent redundancy hampered a detailed investigation of the final steps of NMD so far, since inactivation of one decay route seemed to be partially compensated by the other.

Here we addressed the central question if and how SMG5-SMG7 and SMG6 functionally cooperate and influence each other. We hypothesized that the inactivation of SMG5-SMG7 should activate the SMG6-dependent NMD pathway and still permit normal NMD if both pathways are independent. However, we show here that the combined loss of SMG5-SMG7 completely inactivates NMD. Contrary to our expectations, SMG6 was inactive in cells depleted of SMG5-SMG7, indicating that SMG6 is not independent and could not compensate for the loss of the deadenylation-dependent branch of NMD. This was especially surprising given that SMG6 was previously considered to be the dominant NMD-executing factor. Exploring the potential mechanism, we find that the deadenylation-promoting function of SMG7 can enhance target degradation but is in general dispensable for NMD. We propose a model, in which SMG5 and SMG7 recruitment to hyper-phosphorylated UPF1 acts as an additional licensing step required for SMG6-mediated degradation of the target transcript. This model of two-factor authentication explains the tight control of NMD on regular transcripts, which prevents the untimely access of endonucleolytic decay activities.

## Results

### NMD is impaired in SMG7 knockout cells

Phosphorylation of UPF1 represents a central checkpoint in NMD, which is followed by SMG6-mediated endonucleolytic cleavage and/or SMG5-SMG7-mediated deadenylation of the target transcript (Fig. 1a). We hypothesized that after deactivating the deadenylation-dependent NMD branch, the execution of NMD would rely exclusively on the activity of SMG6. To achieve this goal, we generated SMG7 knockout (KO) Flp-In-T-REx-293 cells and identified three clones lacking the SMG7-specific band in western blot analyses using three different antibodies (Fig. 1b and Extended Data Fig. 1a,b). In all clones, the SMG7 genomic locus contained different frame-shift inducing insertions/deletions, which also resulted in altered splicing of the CRISPR-targeted SMG7 exon. (Extended Data Fig. 1c-e). Phenotypically, the SMG7 KO clones proliferated slower compared to the WT cells, with no apparent decrease in cell survival (Extended Data Fig. 1f,g). These results indicate that the depletion of full-length SMG7 protein impairs cellular fitness, presumably due to reduced NMD capacity. To test if NMD is indeed impaired in SMG7 KO cells, we quantified the levels of two exemplary endogenous NMD targets. SRSF2 is a serine/arginine-rich (SR) splicing factor, which auto-regulates its expression by generating NMD-sensitive splice isoforms of its mRNA^74^. ZFAS1 is a snoRNA host mRNA with a short PTC-containing ORF, which was reported as an NMD target undergoing SMG6-dependent endocleavage^70^. The tested SMG7 KO clones (2 and 34), showed strongly upregulated levels of the ZFAS1 mRNA and the NMD-sensitive SRSF2 isoforms (Fig. 1c). Baseline levels of these NMD substrates were restored by expressing the wild type SMG7 protein from genomically integrated constructs. Importantly, the NMD defect was quantitatively more pronounced in the SMG7 KO cells compared to a siRNA-mediated knockdown (KD) of SMG7 in control cells (Fig. 1d).

**Fig. 1:**
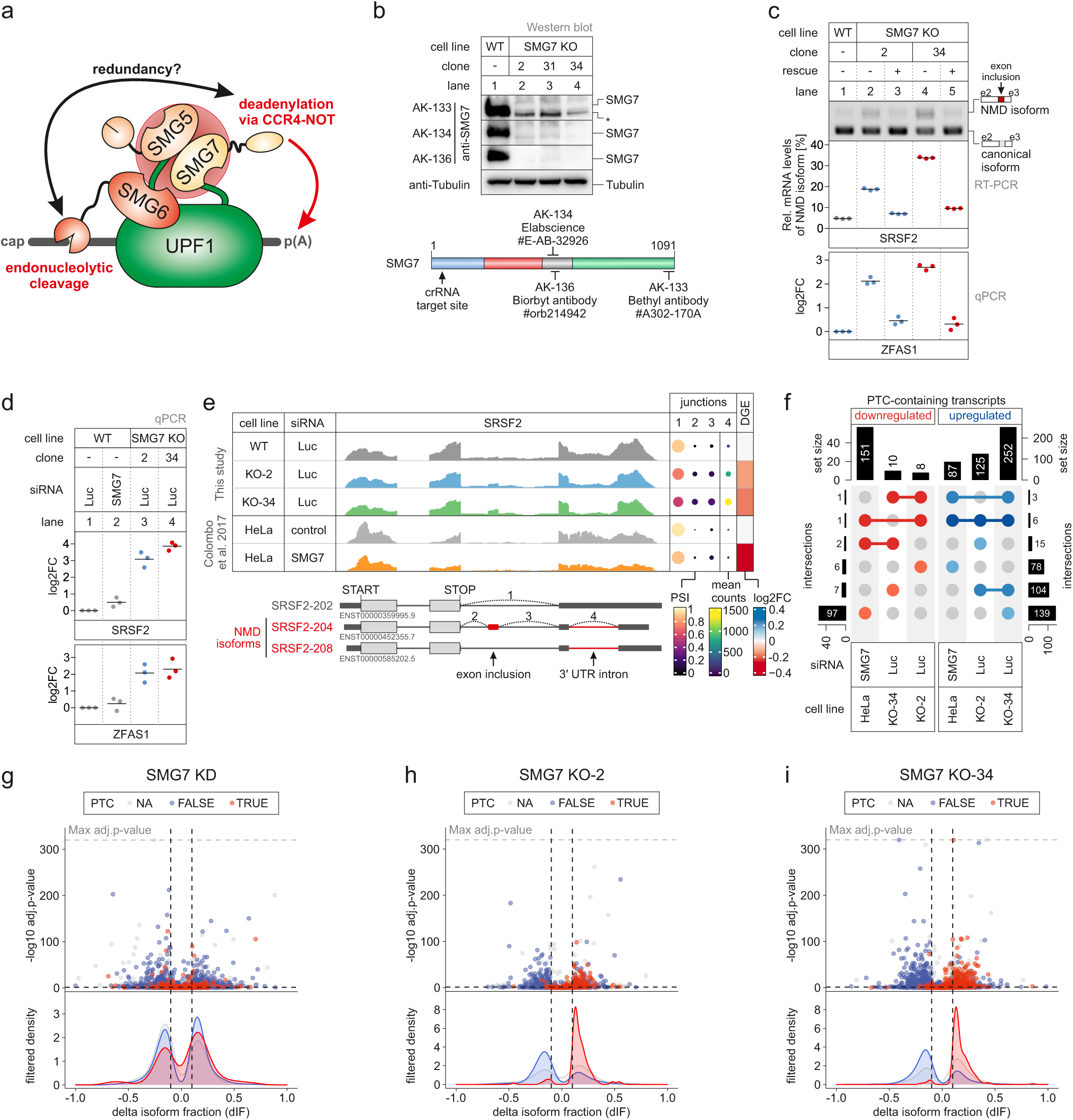
SMG7 depletion impairs NMD activity. **a**, Schematic representation of the final steps of NMD. Phosphorylated UPF1 (indicated by the red sphere) recruits the SMG5-SMG7 heterodimer to the target mRNA, thereby promoting deadenylation. Recruitment of SMG6 to UPF1 results in endonucleolytic cleavage of the target transcript via the SMG6 PIN domain. The SMG5 PIN domain is inactive. **b**, Western blot analysis of SMG7 knockout (KO) cell lines (clones 2, 31 and 34) with the anti-SMG7 antibodies AK-133, AK-134 and AK-136; Tubulin serves as control (see Methods for antibody details). The region of SMG7 detected by the antibodies is schematically depicted and the crRNA targeting site indicated. **c**, End-point RT-PCR detection of SRSF2 transcript isoforms (top) and quantitative RT-PCR-based detection (qPCR; bottom) of ZFAS1 in the indicated cell lines with or without expression of FLAG-tagged SMG7 as rescue construct. The detected SRSF2 isoforms are indicated on the right, the NMD-inducing included exon is marked in red (e = exon). Relative mRNA levels of SRSF2 isoforms were quantified from bands of agarose gels; data points and means from the qPCRs are plotted as log2 fold change (log2FC) (n=3). **d**, Quantitative RT-PCR-based detection (qPCR) of SRSF2 isoforms and ZFAS1 in the indicated cell lines upon treatment with the indicated siRNA. The ratio of NMD isoform to canonical isoform (SRSF2) and ZFAS1 to the C1orf43 reference was calculated; data points and means from the qPCRs are plotted as log2 fold change (log2FC) (n=3). **e**, Read coverage of SRSF2 from SMG7 KO and published SMG7 KD (Colombo et al. 2017) RNA-Seq data is shown as Integrative Genomics Viewer (IGV) snapshots. The canonical and NMD-sensitive isoforms are schematically indicated below. Percent spliced in (PSI; from leafcutter analysis) and mean counts from 4 indicative splice junctions are shown. Differential gene expression (from DESeq2) is depicted as log2 fold change (log2FC) as last column. **f**, Overlap of up- or downregulated premature termination codon (PTC)-containing isoforms between the SMG7 KO or KD RNA-Seq data is shown as UpSet plot. **g-i**, Volcano plots show different SMG7 depletion RNA-Seq analyses. Isoforms containing annotated PTC (red, TRUE), regular stop codons (blue, FALSE) or having no annotated open reading frame (gray, NA) are indicated. The change in isoform fraction (dIF) is plotted against the −log10 adjusted p-value (adj.p-value). Density plots show the distribution of filtered PTC positive/negative isoforms in respect to the dIF, cutoffs were |dIF| > 0.1 and adj.p-value < 0.05.

To gain insights into the transcriptome-wide effects of the SMG7 depletion, we sequenced poly(A)+ enriched mRNA from both SMG7 KO clones and identified differentially expressed genes (Extended Data Fig. 2a and Supplementary Table 1). Consistent with the mRNA-degrading function of SMG7 in NMD, more than twice as many genes were upregulated than downregulated in the SMG7 KO cells (Extended Data Fig. 2b). Compared to a recently published study using SMG7 KD in HeLa cells^14^, this ratio of upregulated vs. downregulated genes was higher (Extended Data Fig. 2c-e). We observed a substantial overlap between upregulated genes in both SMG7 KO cell lines, indicating that these genes are high-confidence SMG7 targets. In contrast, only a limited overlap between downregulated genes could be detected, suggesting these are rather clone-specific effects or off-targets (Extended Data Fig. 2b). From these analyses and the small overall overlap of differentially expressed genes in SMG7 KO cells compared to the SMG7 KD in HeLa cells, we conclude that the complete SMG7 depletion leads to stronger NMD inhibition than the KD.

Next, we quantified alternative splicing events (Supplementary Table 2), as well as differential transcript usage and identified significant isoform switches (Extended Data Fig. 2a and Supplementary Table 3). Isoform switches are characterized by significant changes in the relative contribution of isoforms to the overall gene expression when comparing two conditions^75^. Because of the identification at the isoform level, this approach allows the identification of PTC-containing transcripts that are upregulated upon NMD inhibition. As a specific example of such an isoform switch, we visualized the read coverage for the previously used bona fide NMD target SRSF2. While the overall SRSF2 expression remained nearly unchanged, we detected prominent NMD-inducing exon inclusion and 3′ UTR splicing events in the SMG7 KO but not in the SMG7 KD conditions (Fig. 1e). We verified the SMG7-dependent upregulation of additional examples by end-point PCR (Extended Data Fig. 2f-h).

On a transcriptome-wide scale, NMD-sensitive isoforms with annotated PTC were almost exclusively detected to be upregulated in the SMG7 KO cells, which was not the case in the SMG7 KD in HeLa cells^14^ (Fig. 1f-i). Whereas the functional overlap of isoform switches between the SMG7 KO clones was evident, only a mild enrichment compared to the published SMG7 KD data could be detected (Extended Data Fig. 2i-k). Collectively, the RNA-Seq data analysis supported the initial observation that NMD is robustly impaired when SMG7 is knocked out. Due to the clear effect on NMD upon complete loss of SMG7, the KO cells provide an ideal background to examine further mechanistic details of NMD, which could not be studied before.

### SMG5 is required to maintain residual NMD in SMG7-depleted cells

We utilized the SMG7 KO cells to investigate the molecular features and protein-protein interactions required by SMG7 to support NMD. Specifically, we aimed to confirm whether SMG7 initially binds phosphorylated UPF1 (p-UPF1) and subsequently triggers deadenylation of the target mRNA via the recruitment of the CCR4-NOT complex (Fig. 1a). To this end, we generated stable SMG7 KO clone 2 cell lines that inducibly express SMG7 variants as rescue proteins. Whereas the wild type SMG7 was shown to fully rescue the NMD defect (Fig. 1c), the 14-3-3^mut^ (unable to interact with p-UPF1)^76^ was expected to be inactive (Fig. 2a). However, both wild type and mutant proteins efficiently restored NMD activity in the SMG7 KO cells (Fig.2b). This suggests that the p-UPF1 binding is not absolutely critical for the function of SMG7 in NMD. Next, we investigated if SMG7 has to form a heterodimer with SMG5 for full NMD activity. Surprisingly, the expression of a G100E mutant of SMG7 (unable to interact with SMG5)^62^ (Fig. 2c) failed to rescue the NMD defect (Fig. 2a,b). This finding was unexpected in the light of the currently advocated NMD model, in which SMG5 is merely a companion for SMG7 with the role to potentially strengthen the binding of the SMG5-SMG7 heterodimer to p-UPF1^62^.

**Fig. 2:**
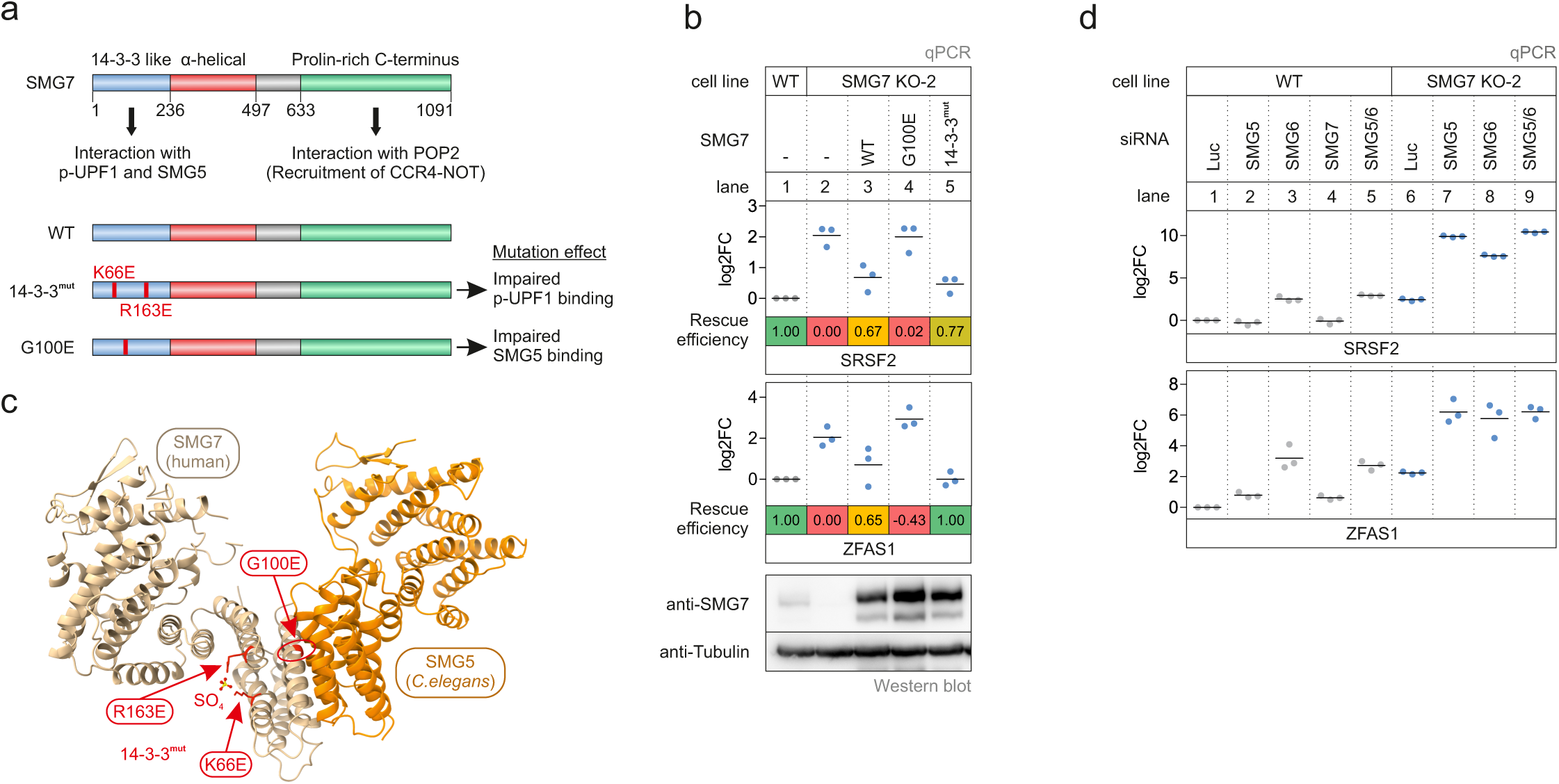
SMG7 requires the interaction with SMG5 to rescue the SMG7 knockout phenotype. **a**, Schematic representation of the SMG7 domain structure. The proposed functions of the domains are indicated and mutated constructs and their expected effect are shown below. **b**, Quantitative RT-PCR-based detection (qPCR) of SRSF2 isoforms and ZFAS1 was carried out in the indicated cell lines upon expression of the indicated FLAG-tagged rescue constructs. The ratio of NMD isoform to canonical isoform (SRSF2) and ZFAS1 to the C1orf43 reference was calculated; data points and means from the qPCRs are plotted as log2 fold change (log2FC) (n=3). Western blot analyses are shown below. Tubulin serves as control. **c**, Model of the SMG5-SMG7 heterodimer structure. Human SMG7 (PDB ID: 1YA0) was modelled on the *C. elegans* SMG5-SMG7 structure (PDB ID: 3ZHE). Critical SMG7 mutations are highlighted in red and the highlighted sulphate ion mimics a phosphorylated residue. **d**, Quantitative RT-PCR-based detection (qPCR) of SRSF2 isoforms and ZFAS1 was carried out in the indicated cell lines upon treatment with the indicated siRNA. The ratio of NMD isoform to canonical isoform (SRSF2) and ZFAS1 to the C1orf43 reference was calculated; data points and means from the qPCRs are plotted as log2 fold change (log2FC) (n=3).

This finding prompted us to systematically address the question, which combinations of the three decay-inducing proteins SMG5, SMG6 and SMG7 are required for NMD (Fig. 2d). Single SMG5 or SMG7 KDs in wild type 293 cells resulted in very mild or nearly undetectable inhibition of NMD (Fig. 2d; lanes 2 and 4), whereas depletion of SMG6 showed an intermediate effect, reflected by the upregulation of the SRSF2 NMD isoform and ZFAS1 (Fig. 2d; lane 3). Co-depletion of SMG6 and SMG5 via siRNAs showed a similar inhibitory effect on NMD as the single SMG6 KD (Fig. 2d; lane 5 vs. lane 3). As expected, KD of SMG6 in the SMG7 KO cells strongly abolished NMD activity, since both endonucleolytic and exonucleolytic pathways of NMD are inactivated (Fig. 2d; lane 8). Remarkably, an even more dramatic NMD inhibition was achieved by depleting SMG5 in the SMG7 KO cells, which could not be further enhanced by the additional KD of SMG6 (Fig. 2d; lane 9 vs. lane 7). This result corroborates the failed rescue with the SMG7 G100E mutant and shows that the SMG5-SMG7 heterodimer is critical for NMD, even when SMG6 is present. Therefore, these observations profoundly question the independence of the SMG5-SMG7 and SMG6 decay pathways and suggest a functional hierarchy in NMD execution.

### NMD is completely inhibited transcriptome-wide upon loss of SMG5-SMG7

The strongly impaired NMD in cells depleted of SMG5 and SMG7 encouraged us to sequence mRNA from SMG7 KO cells (clones 2 and 34) with an additional SMG5 or SMG6 KD (Extended Data Fig. 3a). As expected for complete NMD inhibition, the combined depletion of SMG6 and SMG7 resulted in massive changes of gene expression and isoform usage, which were qualitatively and quantitatively comparable to published SMG6-SMG7 double KD in HeLa cells^14^ (Fig. 3a-c and Extended Data Fig. 3b-d). Whereas SMG5 KD in control cells had very mild effects on the transcriptome, downregulation of SMG5 in SMG7 KO cells exhibited equal or even more pronounced changes in gene expression and isoform usage compared to the SMG6-SMG7-depleted condition (Fig. 3d-f and Extended Data Fig. 3e-g). As a representative example, the alternative splicing pattern of SRSF2 displayed a complete switch from the normal to NMD-sensitive isoforms when SMG5 or SMG6 were depleted in SMG7 KO cells (Fig. 3g). Interestingly, the highest overlap of upregulated NMD-sensitive isoforms was found between both SMG7 KO cell lines with SMG5 or SMG6 KD, suggesting that these four conditions predominantly target the same transcripts (Extended Data Fig. 3h).

**Fig. 3:**
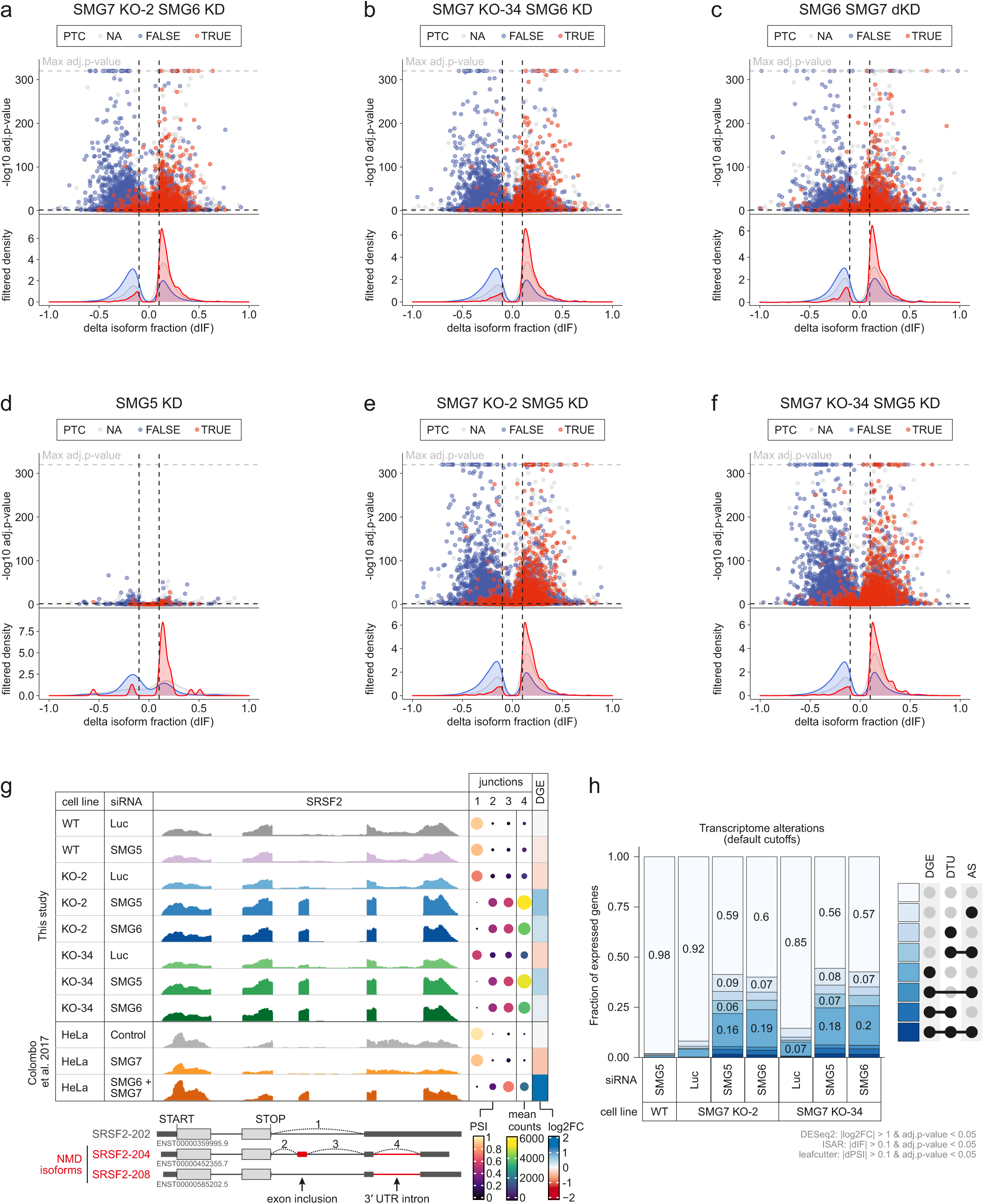
Downregulation of SMG5 in SMG7 knockout cells completely inactivates NMD. **a-f**, Volcano plots show the indicated RNA-Seq analyses. Isoforms containing annotated PTC (red, TRUE), regular stop codons (blue, FALSE) or having not annotated open reading frame (grey, NA) are indicated. The change in isoform fraction (dIF) is plotted against the −log10 adjusted p-value (adj.p-value). Density plots show the distribution of filtered PTC positive/negative isoforms in respect to the dIF, cutoffs were |dIF| > 0.1 and adj.p-value < 0.05. **g**, Read coverage of SRSF2 from SMG7 KO plus knockdown and published knockdown (Colombo et al. 2017) RNA-Seq data is shown as Integrative Genomics Viewer (IGV) snapshots. The canonical and NMD-sensitive isoforms are schematically indicated below. Percent spliced in (PSI; from leafcutter analysis) and mean counts from 4 indicative splice junctions are shown. Differential gene expression (from DESeq2) is depicted as log2 fold change (log2FC) as last column. **h**, Fraction of expressed genes (genes with non-zero counts in DESeq2) which undergo differential gene expression (DGE), differential transcript usage (DTU) and/or alternative splicing (AS) was calculated using the respective computational analysis.

Intrigued by the remarkable changes in gene expression and isoform usage when NMD was completely inhibited, we wanted to re-examine the statement that about 10% of genes are affected by NMD. To this end, we calculated how many genes showed single or combined differential gene expression (DGE), differential transcript usage (DTU) or alternative splicing (AS) events when the SMG7 KO was combined with the KD of SMG5 or SMG6 (Fig. 3h). With this approach, we find that about 40% of the expressed genome in the Flp-In-T-REx-293 cells is under direct or indirect control of NMD. With more stringent cutoffs for the analyses, still around 20% of all expressed genes were affected (Extended Data Fig. 3i), mostly those with medium to high expression levels (Extended Data Fig. 3j). Collectively, the RNA Seq analysis confirmed that SMG5 KD, as well as SMG6 KD, have similar effects on NMD in SMG7 KO cells. It also provided global evidence that the loss of the SMG5-SMG7 heterodimer completely inactivates NMD and leads to massive changes in the expressed transcriptome.

### Loss of SMG5 and SMG7 prohibits endonucleolytic cleavage of NMD substrates

The observed NMD inhibition upon the co-depletion of SMG5 and SMG7 suggests that SMG6 is equally inactivated, although it is widely assumed that SMG6 acts redundantly to and independently of SMG5-SMG7^14,64^. This unexpected result raised the question, whether SMG6 requires the presence of SMG5-SMG7 for endonucleolytic cleavage of its target mRNAs during NMD. We monitored the activity of SMG6 by northern blotting, which allows us to detect decay intermediates resulting from endonucleolytic or 5′-3′ exonucleolytic degradation. To this end, stably integrated triosephosphate isomerase (TPI) mRNA reporters were expressed as wild type or NMD-inducing PTC160 variant in control or SMG7 KO cells with different combinations of siRNA-mediated knockdowns (Fig. 4a). The reporter mRNA also contained XRN1-resistant sequences (xrRNAs) in the 3′ UTR that produce meta-stable intermediates of 5′-3′ decay (called xrFrag) and thereby provided information about the extent and directionality of mRNA degradation^72,77^. Upon depletion of the major cytoplasmic 5′-3′ exonuclease XRN1, SMG6-generated endonucleolytic cleavage products (designated 3′ fragments) of the PTC-containing reporter mRNA are detected as an additional band (Extended Data Fig. 4a; lane 6). Of note, the isolated SMG7 KO or SMG6 KD resulted in a slight accumulation of the full-length reporter (Extended Data Fig. 4a; lanes 8 and 10), indicating partial NMD inhibition consistent with the literature and our earlier observations. However, we did not observe increased 3′ fragment levels upon loss of SMG7, which indicates that the SMG6 activity is not compensatory in the absence of SMG7 (Fig. 4b and Extended Data Fig. 4a,b; lane 14). While KD of SMG5 in control cells had no inhibitory effect on endonucleolytic cleavage, depletion of SMG5 in SMG7 KO cells completely abolished the generation of 3′ fragments (Fig. 4b and Extended Data Fig. 4b; lanes 4 and 8 vs. 12 and 16). Furthermore, the accumulation of the PTC160 reporter mRNA to WT levels and the disappearance of the xrFrag band confirmed that NMD is completely inactivated in SMG5-SMG7-depleted conditions. The dramatic effect on the endonucleolytic cleavage activity was further confirmed by investigating the naturally occurring stable cleavage product of NOP56 (Fig. 4c)^78^. In the control cells, the cleavage product was predominantly present, but became undetectable in the SMG5-SMG7-depleted condition. Taken together, these results underline the previous observation that the SMG5-SMG7 heterodimer is required for general NMD activity and, surprisingly, also for SMG6 activity.

**Fig. 4:**
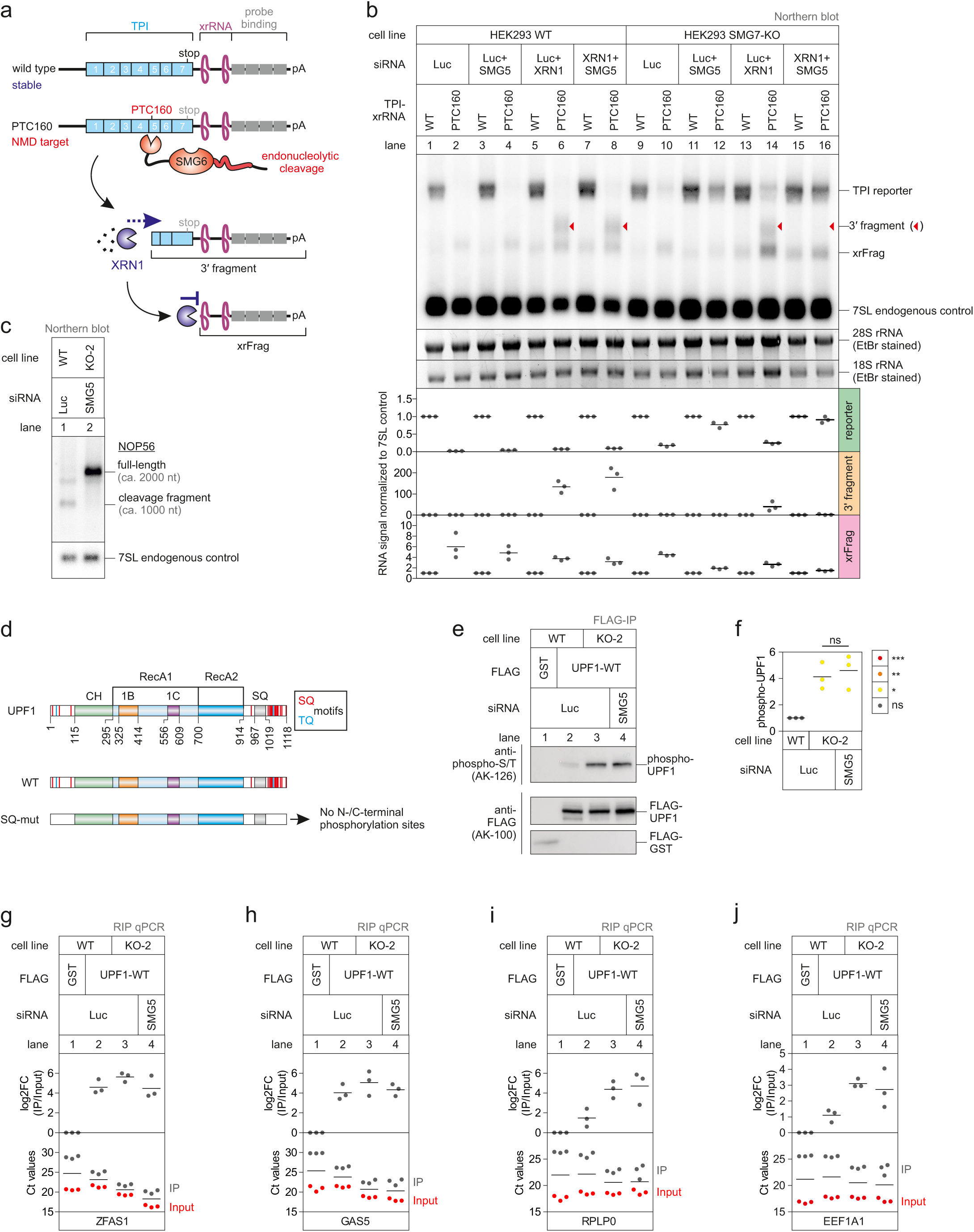
SMG6 endonucleolytic cleavage is inactivated in SMG5-SMG7 depleted cells. **a**, Schematic overview of the triosephosphate isomerase (TPI) reporter constructs and their functional elements. **b**, Northern blot analysis of TPI reporter, 3′ fragments, xrFrag and 7SL endogenous control. Ethidium bromide stained 28S and 18S rRNAs are shown as additional controls. Quantification results are shown as data points and mean (n=3). **c**, Northern blot analysis of endogenous NOP56. Different transcript isoforms are indicated. **d**, Schematic overview of UPF1, highlighting the putative SQ/TQ phosphorylation sites. **e-f**, Analysis of UPF1 phosphorylation status after IP of expressed FLAG-tagged UPF1 (**f**) with quantification (**g**). Quantification results are shown as data points and mean (n=3). **g-j**, qPCR detection of ZFAS1 (**h**), GAS5 (**i**), RPLP0 (**j**) and EEF1A1 (**k**) was carried out in IP and Input samples after RNA immunoprecipitation (RIP) with FLAG-tagged UPF1. Data points and means from the qPCRs are plotted as log2 fold change (log2FC) (n=3). Raw Ct values are shown for comparison.

### UPF1 phosphorylation and target-discrimination is altered upon SMG7 knockout

The common denominator of SMG5-SMG7 and SMG6 in the NMD pathway is UPF1 serving as the binding platform on the target mRNA. Controlled phosphorylation cycles of UPF1 are required for functional NMD, since disturbed UPF1 dephosphorylation inhibits NMD^52,53,60^. While SMG5 and SMG7 exclusively bind to phosphorylated UPF1^53,60,62^, SMG6 interacts with UPF1 also in a phosphorylation-independent manner^63,79^. Intriguingly, the p-UPF1 binding proteins SMG5 and SMG7 were implicated in the dephosphorylation of UPF1, potentially via the recruitment of the protein phosphatase 2A (PP2A)^53,61,80,81^. Therefore, we wanted to explore the possibility, that an altered UPF1 phosphorylation status is responsible for the NMD inhibition in SMG5-SMG7-depleted cells. To this end, we expressed FLAG-tagged UPF1 wild type or a mutant lacking all N- and C-terminal potential phosphorylation sites (SQ-mut) in wild type or SMG7 KO cells (Fig. 4d). Wild type UPF1, but not the SQ-mutant, showed increased phosphorylation in the SMG7 KO cells, which did not increase further when SMG5 was depleted (Fig. 4e,f and Extended Data Fig. 5a). However, the overall amount of p-UPF1 in the SMG7 KO cells was relatively small in comparison to cells treated with okadaic acid, a potent inhibitor of protein phosphatase 2A (PP2A) (Extended Data Fig. 5a)^52^. These results confirm that SMG7 is involved in the dephosphorylation of UPF1. However, it seems unlikely that the accumulation of hyperphosphorylated UPF1 represents the main reason for the complete inhibition of NMD and SMG6 activity in SMG5-SMG7 depleted cells.

Previous publications reported that SMG5 and SMG7 stabilise p-UPF1 binding to target mRNAs^61^ and ATPase-deficient UPF1 mutants accumulate in a hyper-phosphorylated, SMG5-SMG7 bound state^48^. Intrigued by these reports, we next investigated whether the NMD target binding or recognition ability of UPF1 is impaired in SMG5-SMG7 depleted cells. To this end, we employed UPF1 RNA immunoprecipitation (RIP) assays to study NMD target discrimination by UPF1^48^. Binding of UPF1 to two NMD targets, which displayed increased mRNA levels upon NMD inhibition (Extended Data Fig. 5b,c), remained unchanged in control, SMG7 KO, or SMG7 KO plus SMG5 KD conditions (Fig. 4g,h). These results suggest that UPF1 is still able to identify and bind to NMD-targeted transcripts, although their degradation cannot be executed anymore. In contrast, non-NMD targets such as RLP0 or EEF1A1, which are normally weakly or not at all bound by WT UPF1 (Extended Data Fig. 5d,e), exhibited stronger association with UPF1 in SMG7 KO cells (Fig. 4i,j). Of note, the impaired release of UPF1 from non-NMD targets resembles the behaviour of ATPase-deficient UPF1 (Extended Data Fig. 5d,e). Although these results suggest that the loss of SMG7 impairs the target discrimination ability of UPF1, the additional KD of SMG5 did not substantially increase this effect (Fig. 4i,j). In summary, the depletion of SMG7 results in dysregulation of the UPF1 phosphorylation status and target discrimination, while no convincing evidence for additive effects of SMG5 knockdown on these defects were found.

### SMG7-mediated deadenylation is dispensable for NMD

To gain mechanistic insight into the requirements of SMG5 and SMG7 for functional NMD, we repeated the SMG7 rescue experiments with additional combinations of rescue constructs and KDs of SMG5 or SMG6 (Fig. 5a). Consistent with our earlier observations, SMG7 WT rescued the SMG7 defect in all conditions, whereas a 14-3-3^mut^ / G100E double-mutation completely inactivated SMG7 in the full-length context (Fig. 5b; lane 3 vs. lane 6). We were surprised that a SMG7 1-633 deletion mutant (unable to recruit the CCR4-NOT complex)^64^ efficiently rescued NMD in SMG7 KO cells, even when SMG5 or SMG6 were depleted in addition (Fig. 5b; lane 7). This result indicates that the deadenylation of NMD substrates by SMG7 is not required for their efficient degradation. In contrast to a previous report^64^, the C-terminus of SMG7 is not required for NMD even when SMG6 is downregulated. However, by combining the C-terminal truncation with either 14-3-3^mut^ or G100E mutations, these SMG7 variants became less NMD-competent, especially in the SMG5 KD condition (Fig. 5b; lanes 4-5 vs. lanes 8-9). This observation suggests that the deadenylation-inducing activity might serve as an additive but dispensable feature that helps to clear NMD targets, especially when other features of SMG7 are inactivated.

**Fig. 5:**
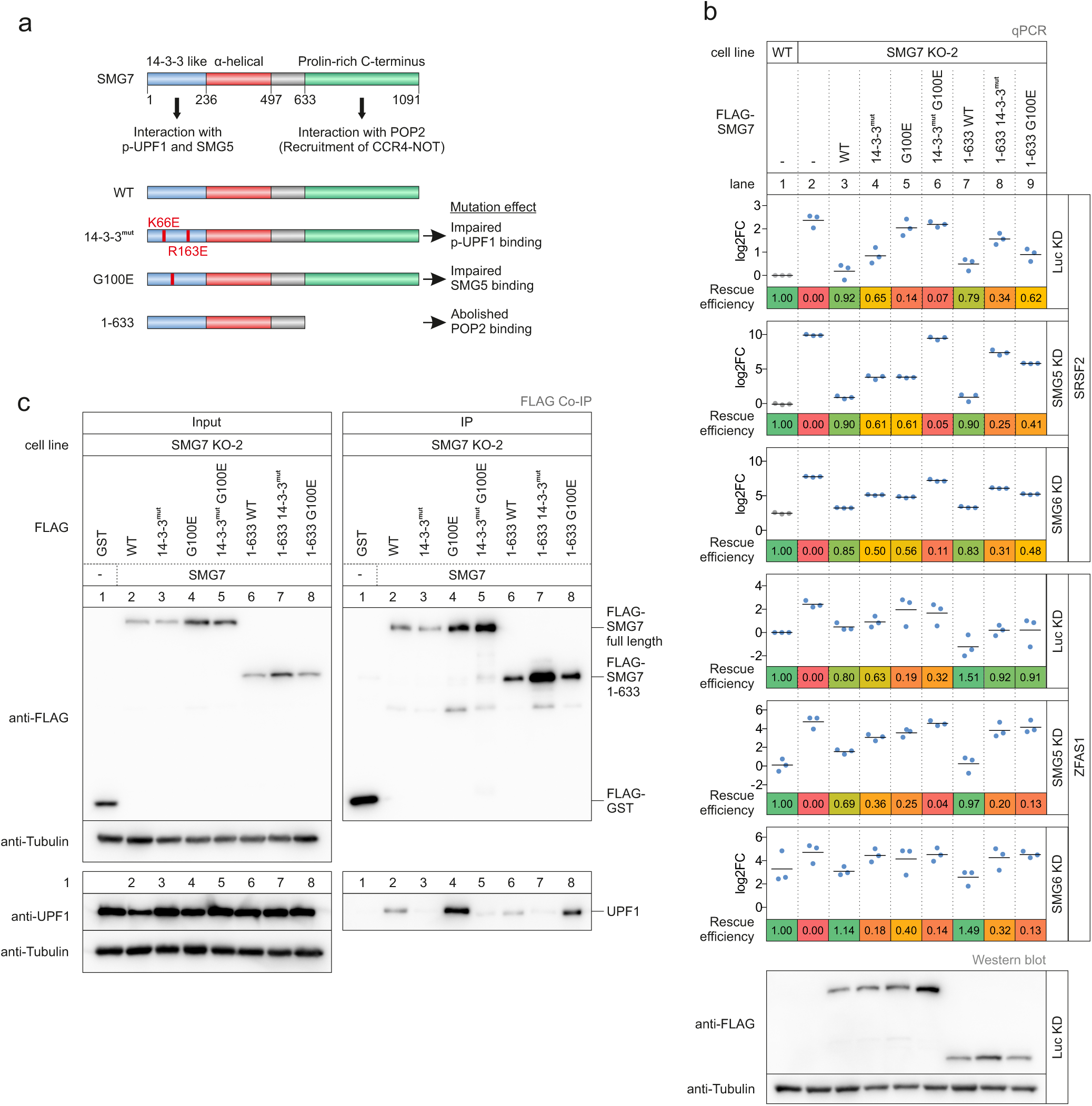
SMG7 supports NMD independently of the deadenylation-promoting function. **a**, Schematic representation of the SMG7 domain structure. The proposed functions of the domains are indicated and mutated constructs and their expected effect are shown below. **b**, Quantitative RT-PCR-based detection (qPCR) of SRSF2 isoforms and ZFAS1 was carried out in the indicated cell lines upon expression of the indicated FLAG-tagged rescue constructs and knockdown with the indicated siRNA. The ratio of NMD isoform to canonical isoform was calculated; data points and means from the qPCRs are plotted as log2 fold change (log2FC) (n=3). Western blot analyses are shown below. Tubulin serves as control. **c**, Western blot after co-immunoprecipitation of FLAG-tagged GST (control) or SMG7 constructs. Tubulin serves as control.

Next, we performed co-IP experiments with the functionally tested set of FLAG-tagged SMG7 constructs to analyse their steady-state interaction with UPF1 (Fig. 5c). The full-length or truncated 14-3-3^mut^ protein was unable to co-IP UPF1, consistent with the role of the 14-3-3 like domain to mediate the interaction with phosphorylated UPF1 (Fig. 5c; lane 2 vs. lanes 3 and 7)^63,79^. In turn, this result also indicates that SMG5 does not bridge the 14-3-3^mut^ SMG7 protein to UPF1. Interestingly, the G100E mutants co-immunoprecipitated more UPF1 than the respective wild type construct (Fig. 5c; lanes 2,6 vs lanes 4,8). This finding confirms the assumption that SMG7 does not require the heterodimerization with SMG5 to interact with UPF1^62^. In conjunction with the failed functional rescue of the G100E mutants, this result implies that despite prominent UPF1-SMG7 binding, the interaction of SMG7 with SMG5 might be important to advance in the NMD process.

### SMG5 functionally complements the loss of SMG7

These unexpected observations concerning SMG7 prompted us to investigate the molecular properties of SMG5. Specifically, we aimed to identify which function of SMG5 is required to support NMD in the absence of SMG7 (Fig. 6a). The first striking result was the almost complete rescue of the SMG7 depletion phenotype by the expression of SMG5 WT or G120E mutant (unable to interact with SMG7)^62^ in control or SMG5 KD conditions (Fig. 6b; lanes 3 and 5). This finding was very surprising, given that SMG5 has no reported ability to support NMD directly. Although SMG5 could potentially execute endonucleolytic cleavage via the PilT N-terminus (PIN) domain that structurally resembles the functional PIN domain of SMG6, two of the three required catalytic residues are missing in the inactive C-terminal SMG5 PIN domain (Fig. 6a)^82^. SMG5 was also shown in initial reports to directly recruit decapping factors, which could explain the NMD activity we observe in the rescue assays^83^. However, later studies could not confirm this path of SMG5-dependent decapping and rather pointed to UPF1 directly interacting with general decapping factors^64,65^.

**Fig. 6:**
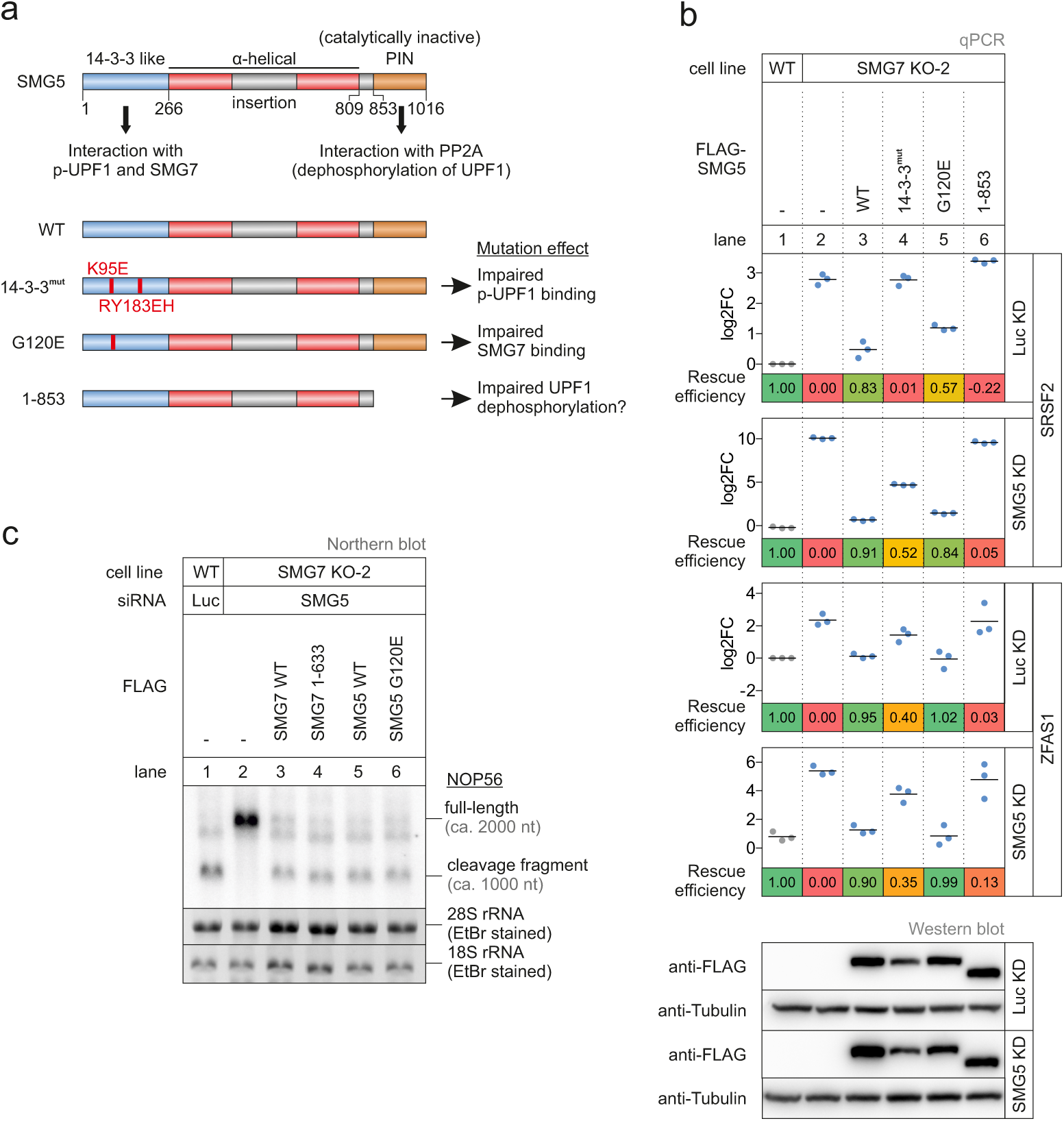
SMG5 expression rescues the SMG7 KO phenotype. **a**, Schematic representation of the SMG5 domain structure. The proposed functions of the domains are indicated and mutated constructs and their expected effect are shown below. **b**, Quantitative RT-PCR-based detection (qPCR) of SRSF2 isoforms and ZFAS1 was carried out in the indicated cell lines upon expression of the indicated FLAG-tagged rescue constructs and knockdown with the indicated siRNA. The ratio of NMD isoform to canonical isoform was calculated; data points and means from the qPCRs are plotted as log2 fold change (log2FC) (n=3). Western blot analyses are shown below. Tubulin serves as control. **c**, Northern blot analysis of endogenous NOP56 (n=3). Different transcript isoforms are indicated.

Altogether, the observation of SMG5 expression rescuing the loss of SMG7 once more questions the relevance of the SMG7 deadenylation-promoting function for NMD. We hypothesized that in the absence of SMG7, SMG5 might rely on its N-terminal 14-3-3-like domain to interact with p-UPF1 to activate NMD. In line with this hypothesis, mutating three residues in the potential phosphopeptide binding pocket of SMG5 severely affected the ability of SMG5 to rescue the SMG7 KO phenotype in control or SMG5 KD conditions (Fig. 6b; lane 4). This result indicates that either the interaction between SMG5 and UPF1 per se suffices to rescue NMD or once this interaction is established, another function of SMG5 is required to maintain NMD capacity. To test these ideas, we generated a SMG5 deletion mutant lacking the C-terminal PIN domain (Fig. 6a), which should still be able to interact with UPF1. This SMG5 construct comprising the first 853 amino acids was unable to restore NMD activity (Fig. 6b; lane 6), suggesting that the catalytically inactive PIN domain of SMG5 is essential to support NMD. In search of an explanation, we considered early reports that the C-terminus of SMG5 might be involved in the dephosphorylation of UPF1, likely by direct recruitment of the PP2A complex^53,60^. Although we did not observe further hyper-phosphorylation of UPF1 in the SMG7 KO plus SMG5 KD cells (Fig. 4e,f), we cannot exclude a function of SMG5 in the process of UPF1 dephosphorylation. Conceivably, the residual SMG5 proteins left after the KD are sufficient to counteract further UPF1 hyper-phosphorylation, whereas NMD cannot be supported anymore.

Finally, we were intrigued by the fact that both SMG5 and SMG7 wildtype proteins can individually rescue the SMG7 KO phenotype. Specifically, we wondered if the main NMD-supporting function of both factors could be to enable SMG6-mediated endonucleolytic cleavage of the target mRNA. To test this hypothesis, we used the NOP56 cleavage product as an indicator for SMG6 activity, since the generation of this meta-stable RNA fragment was shown to be strongly SMG6-dependent^78^. All SMG5 and SMG7 rescue proteins that restored full NMD activity also resulted in normal NOP56 cleavage pattern, indicating that SMG6 was reactivated in these cells (Fig. 6c). Based on these results, we postulate that SMG5 and SMG7 maintain NMD competence by permitting the activation of SMG6.

## Discussion

The correct execution of NMD not only prevents the production of aberrant gene products, but also shapes the transcriptome on a global scale^43^. NMD is generally perceived as a robust, but highly dynamic process that integrates different inputs, including mRNP composition and translational status, in order to efficiently identify and remove transcripts that appear to be faulty^42^. Multiple RNA degradation pathways can be employed after identification of target transcripts, which are all centred around the key factor UPF1 and provide reliable elimination of the mRNA. UPF1 can directly or indirectly interact with the general mRNA decapping complex^64,83–87^, which could enhance the degradative power of NMD, but the major decay paths during NMD utilize the UPF1-recruited SMG5-SMG7 and SMG6. Although evidence pointed to the independence of these branches in the past, we show here that SMG6 cannot endonucleolytically cleave NMD substrates in cells lacking the SMG5-SMG7 heterodimer (Fig. 4 and Extended Data Fig. 4), resulting in complete NMD inactivation. Therefore, functional dependence and hierarchy exists between both pathways.

The reason why this dependency has not been detected so far has probably technical reasons. Virtually all previous experiments that address the interplay between SMG5, SMG6 and SMG7 utilized individual or combined gene silencing of NMD factors depending on siRNA- or shRNA-mediated knockdown. As reported before, the downregulation of SMG5 and/or SMG7 by knockdowns only slightly impairs NMD^35,62^. We show here that a complete and sustained depletion of SMG7 is needed to detect a considerable NMD defect (Fig. 1 and Extended Data Fig. 1). This condition is not achieved by SMG7 knockdown approaches. Furthermore, the downregulation of SMG5 substantially affects NMD activity only in the SMG7 KO conditions (Fig. 3 and Extended Data Fig. 3). Therefore, we propose that a conventional downregulation of the SMG5-SMG7 heterodimer is not sufficient to abolish its function. It was reported before that SMG5 and SMG7 form stable and long-living complexes^35^ and residual heterodimers could potentially outlive the experimental timeframe of knockdown experiments. Remarkably, strongly reduced levels of the SMG5-SMG7 heterodimer after knockdown are still able to support NMD, although both proteins are about two orders of magnitude less abundant than UPF1^88,89^. This observation indicates that the basal levels of SMG5 and SMG7 provide enough buffer capacity to tolerate the partial loss of individual NMD factors or to cope with increasing amounts of NMD targets, e.g., resulting from reduced transcriptomic fidelity. In line with this idea, previous attempts to “overload” the NMD machinery by transiently overexpressing large quantities of NMD substrates did not result in reduced NMD activity^90^.

The remarkable capacity of the NMD process is also reflected in the amount of differentially regulated transcripts that accumulate in cells with inactive NMD. Earlier studies estimated that about 5-10 % of all human genes are directly or indirectly influenced by NMD^9,12–15^. If we consider gene- and isoform-specific effects (differential gene expression, isoform switches, alternative splicing), we find that between 20 and 40 % of the expressed genes are affected by NMD. These values are considerably higher than previous estimates, which can be partially explained by using state-of-the-art RNA-sequencing methods and recent bioinformatic algorithms, allowing a more thorough analysis of the transcriptomic alterations. Furthermore, we believe that the SMG7 KO cells in combination with SMG5 or SMG6 KD result a more efficient NMD inhibition, which could not be achieved with previous attempts based on RNA interference alone. Admittedly, not all of the detected changes in NMD-incompetent cells are direct effects of NMD inhibition, since the misregulation of targets such as the splicing factor SRSF2 will undoubtedly cause secondary effects on the transcriptome. However, the large number of NMD-regulated genes can explain why NMD is essential for cell survival, proliferation and differentiation. It is difficult to imagine that important and fundamental biological processes can function normally without being influenced by the 20-40 % NMD-regulated genes. Given this large amount of potential cellular NMD substrates, it will be important in the future to identify and characterize which mRNA isoforms are authentic NMD-regulated transcripts. This will also help to understand the process of NMD better and establish further rules for NMD activating features^91^.

On a conceptional level, NMD identifies and degrades transcripts that fail to pass quality control standards. To this end, the NMD pathway makes use of a multitude of potent cytoplasmic RNA degradation tools, such as the endonuclease SMG6. However, the access to and application of these tools must be very tightly controlled to minimise spurious degradation of normal transcripts. This control is especially important, since NMD probably monitors every translation termination event and uncontrolled SMG6-mediated mRNA cleavage would be catastrophic for the cell. Based on the intensive previous work of several different labs and the data presented in this study, we propose an improved model for the activation of NMD with the ambition to integrate and explain as many earlier observations as possible. In our opinion, this can best be accomplished with a two-factor authentication model (Fig. 7). In this model, UPF1 serves as a general surveillance factor, which has to successfully pass at least two different consecutive authentication procedures on true NMD targets to gain access to SMG6-mediated activity.

**Fig. 7:**
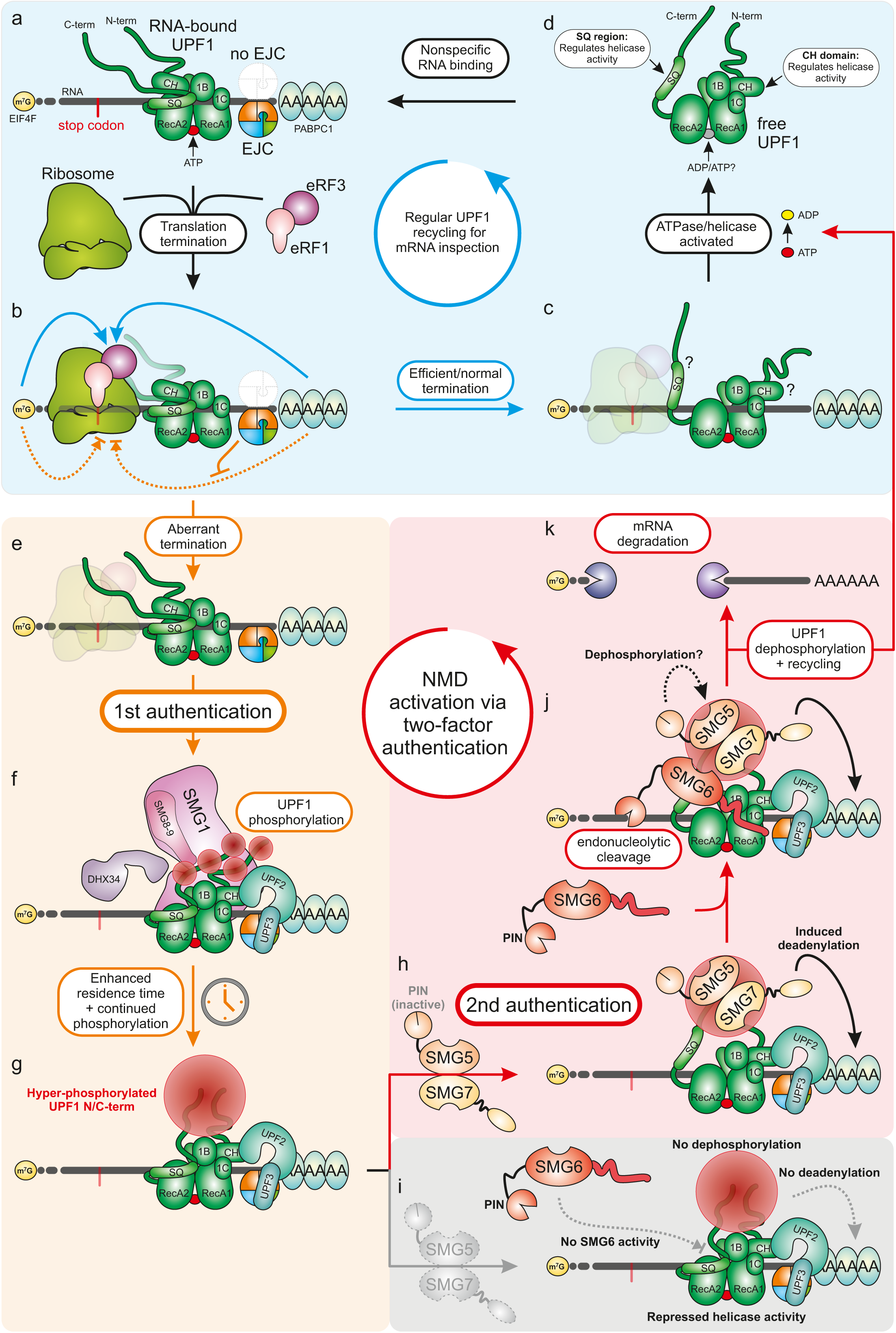
Two-factor authentication model of NMD. **a-d**, Regular inspection cycle of UPF1. UPF1 binds mRNA unspecifically and remains attached until being displaced by the ribosome. UPF1 helicase and ATPase activity is stimulated when ribosomes terminate translation efficiently. Subsequently, UPF1 dissociates from the mRNA and can engage in a new cycle of inspection. **e-g**, First NMD authentication step. Upon aberrant translation termination (e.g. EJC downstream of terminating ribosome), UPF1 remains bound on the mRNA. If an NMD activating features exist, UPF1 is progressively phosphorylated by the kinase SMG1. Continued presence of UPF1 on the mRNA and NMD-activating features lead to hyper-phosphorylated UPF1. **h-k**, Second NMD authentication step. If SMG5-SMG7 are absent, the hyper-phosphorylated UPF1 accumulates and NMD cannot successfully eliminate the target transcript. Also, UPF1 is not recycled. If SMG5-SMG7 are present, the interaction with UPF1 allows the binding of SMG6 to UPF1 in order to catalyse the endonucleolytic cleavage of the target transcript. The target transcript is fully degraded by exonucleolytic activities. SMG5 may subsequently dephosphorylated UPF1, the helicase activity is stimulated and UPF1 is effectively recycled for another round of inspection.

By analogy, we envision that UPF1 acts as a molecular inspector, which is routinely assigned to control the quality of all “end-of-line” transcripts that are translated (Fig. 7a). It has become increasingly clear that the mere binding of UPF1 to an mRNA does not trigger its degradation. This would be in agreement with previous informal names for UPF1 that signify its main function, such as “time bomb” or “kinetic/molecular clock”^4,59,92^. These names indicate that the residence time of UPF1 on a given mRNA influences NMD activity. Accordingly, UPF1 has to be removed from the transcript in a timely manner to prevent NMD in the first place. Towards this goal, translating ribosomes remove most of the RNA-bound UPF1 and only those in the 3’ UTR remain attached to the transcript^44–47^. Furthermore, efficient and proper translation termination probably leads to the activation of the UPF1 ATPase and helicase function, resulting in the dissociation of UPF1 from the transcript^48^ (Fig. 7b-d). The precise molecular mechanism for this termination-dependent UPF1 activation is still largely enigmatic. Either the ribosome itself, the translation termination factors or certain proteins located in the 3’ UTR such as cytoplasmic poly-A binding protein 1 (PABPC1) and polypyrimidine tract binding protein 1 (PTBP1) may contribute to release UPF1 from authentic mRNAs^93–96^. Therefore, proper usage of the transcript for protein production sufficiently convinces the inspector UPF1 that the mRNA is legitimate, resulting in UPF1 being released for another round of inspection (Fig. 7a-d).

When translation terminates inefficiently or at aberrant positions (e.g., upstream of an EJC), the residence time of UPF1 increases, probably due to the lack of strong UPF1 ATPase/helicase activation (Fig. 7b,e). At this point, the first authentication step is installed, which is the progressive SMG1-mediated phosphorylation of UPF1 (Fig. 7f,g). It has been shown that SMG1 associates with and phosphorylates UPF1 preferentially in the presence of UPF2^50,52,57,58,97^, which in turn also alters the conformation and helicase activity of UPF1 upon binding^48,98^. The presence of the EJC and UPF3, which binds to the EJC, is thought to further enhance the kinase activity of SMG1^50^. Finally, SMG1-mediated UPF1 phosphorylation is also controlled by DHX34, SMG8, SMG9 and potentially other auxiliary NMD factors^54–56,58,99^. All these regulatory mechanisms ensure that UPF1 is only phosphorylated when bound to positions on the mRNA, where an NMD-activating arrangement of mRNP features is present. Effectively, the inspector (UPF1) remains attached to the transcript and persisting NMD-activating features will cumulatively lead to hyper-phosphorylated UPF1 (Fig. 7g). To stick to our previous theme, the inspector gets increasingly suspicious that the bound mRNA may be defective.

Notably, earlier reports described that specific phosphorylated residues in the UPF1 N- and C-terminus are critical binding sites for SMG6 and SMG5-SMG7, respectively^50,60,76^. According to this rather rigid concept, degradation could be efficiently induced by even a single UPF1 phosphorylation event. In particular, this is problematic with respect to SMG6, whose recruitment to a barely phosphorylated UPF1 could lead to an untimely endonucleolytic cleavage that cannot be undone. Therefore, we favour the recently emerging view that gradually increasing UPF1 phosphorylation at multiple sites increases the chance for NMD execution^59^.

The combination of UPF1 residence time on a given transcript and phosphorylation status allows for a graduated NMD response. Strong NMD activating features, such as downstream EJCs, efficiently activate SMG1, leading to hyper-phosphorylated UPF1 within a short time frame. In contrast, transcripts with weaker NMD-activating features, such as long 3’ UTRs, likely require longer time and/or translation cycles to accumulate moderately phosphorylated UPF1. Conceptionally, this graduated response allows the rapid degradation of true NMD targets (e.g., PTC-containing transcripts), whereas regular transcripts can be regulated in their expression with varying intensity. Therefore, the first authentication step of NMD does not follow a simple “on/off” scheme, but rather allows for large regulatory flexibility, as was recently proposed^42^.

How is the increasing phosphorylation ultimately converted to an appropriate mRNA degradation response? We propose, based on the existing literature and our data, that a second authentication procedure must be passed to gain access to and/or activate SMG6 (Fig. 7h). Accordingly, we envision an interaction of SMG5-SMG7 with phosphorylated UPF1 as an essential step to permit SMG6-mediated endonucleolytic cleavage of the target mRNA. The evidence supporting this hypothesis is foremost the complete NMD inhibition in SMG5-SMG7-depleted cells, with no detectable SMG6-activity (Fig. 7i). Furthermore, the transcriptomic changes are highly similar between loss of the SMG5-SMG7 heterodimer and the combined SMG7 KO with SMG6 KD, indicating the same functional outcome. This observation is in good agreement with the previously reported extensive redundancy between SMG6 and SMG7^14^. Further support for this model comes from the observation that hyper-phosphorylation alone is not sufficient to induce NMD, as ATPase-deficient UPF1 mutants are phosphorylated but do not support NMD^42,50,59^. Accordingly, we observed increased UPF1 phosphorylation in SMG7 KO cells, despite NMD being impaired.

According to this idea of a second authentication step, SMG6 would remain inactive until SMG5-SMG7 sensed that UPF1 is sufficiently phosphorylated (Fig. 7i). This poses the question, how the access of SMG6 to UPF1 can be controlled? Although SMG6 contains a 14-3-3-like domain and was proposed to interact directly with UPF1 via phosphorylation-dependent binding^60^, a phosphorylation-independent interaction between SMG6 and UPF1 was reported afterwards^63,79,100^. If this phosphorylation-independent interaction were sufficient to activate SMG6-mediated endonucleolytic cleavage, uncontrolled NMD would be observed. Therefore, we propose that the phosphorylation-dependent interaction of SMG5-SMG7 with UPF1 may lead to conformational changes of UPF1, allowing the transient binding and activity of SMG6 on *de facto* authenticated NMD targets (Fig. 7j). The key to this process could be the spatial arrangements of the accessory domains of UPF1. Especially the C-terminally located SQ-region is of interest, as this region was shown to intra-molecularly interact with the helicase domains of UPF1, leading to repression of helicase activity^101^. Intriguingly, this SQ-region, as well as the helicase domain, were reported to mediate the phosphorylation-independent interaction between UPF1 and SMG6^63,79^. It is an attractive hypothesis that SMG5-SMG7-induced dissociation of this SQ-region from the helicase core not only allows the execution of endonucleolytic cleavage by SMG6, but also initiates the ATPase-driven removal of UPF1 from the target. This mode of action would ensure that correctly completed NMD also releases UPF1 from the now partially degraded transcripts, which is required to recycle UPF1 and permit XRN1-mediated elimination of the NMD target (Fig. 7k)^102^. On the other hand, this complex sequence of events would ensure that UPF1 remains locked on transcripts that require further inspection until a decision has been made to either release the inspector UPF1 or to degrade the mRNA. Therefore, in the absence of SMG5-SMG7, phosphorylated UPF1 molecules would accumulate on RNA, being unable to dissociate from the mRNA or to initiate target degradation. This interpretation fits with our UPF1-RIP results and represents an intriguing difference compared to ATPase-deficient UPF1, which was reported to degrade one target molecule, but simply cannot leave this target^102^.

This model undoubtedly raises questions about the molecular function of SMG5 and SMG7, the need to form a SMG5-SMG7 heterodimer, and the mode of SMG5-SMG7 interaction with p-UPF1. Our detailed analysis of SMG5 and SMG7 revealed interesting and unexpected insights into the function of these two proteins. Most impressively, the two proteins seem to exhibit a certain redundancy, because SMG5 or SMG7, respectively rescue the combined SMG5-SMG7 depletion. However, multiple lines of evidence in the past pointed to SMG7 as the dominant and functional component of the SMG5-SMG7 heterodimer. For example, SMG7 can interact with UPF1 even when SMG5 is downregulated or when the binding to SMG5 is impaired (e.g., by the G100E mutant; confirmed in Figure 5C)^62^. Furthermore, SMG7 contributes to the turnover of the target mRNA by recruiting the CCR4-NOT complex^64^. In contrast, the results of our rescue experiments question this SMG7-centered, deadenylation-dependent model. The 1-633 SMG7 deletion mutant, which lacks the CCR4-NOT interaction, could efficiently rescue the SMG7 KO phenotype, even when SMG5 was co-depleted. Therefore, and per our model, the main function of SMG5-SMG7 cannot simply be the recruitment of deadenylation-promoting factors. This interpretation is in line with our result that SMG5 alone can replace SMG7, although SMG5 neither possesses a functional PIN domain^82^ nor interacts with deadenylating complexes. Also, SMG7 1-633 rescues as efficiently as the full-length SMG7 protein when SMG6 is co-depleted, which does not fit with the hypothesis that the SMG7 C-terminus functions redundantly with SMG6^64^. Remarkably, the failed functional rescue, but increased amount of co-immunoprecipitated UPF1 with the SMG7 G100E mutant suggests that strong SMG7-UPF1 binding does not necessarily enhance NMD and that SMG5 plays a more important role than previously anticipated.

What could be the molecular function of SMG5 in NMD? Previous data showed that despite having a 14-3-3-like domain, SMG5 cannot stably interact with p-UPF1 in the absence of SMG7^62^ and SMG5 cannot use the C-terminal PIN domain to eliminate transcripts due to missing catalytic residues^82^. This raises the question, how SMG5 expression is able to rescue the SMG7 KO (Fig. 6). One important aspect that has not been covered in the above-discussed model is the dephosphorylation of UPF1. After completion of NMD, hyper-phosphorylated UPF1 will be present either in free or RNA-bound form. Before this p-UPF1 can be reused for another inspection cycle, it has to be dephosphorylated. Evidence from *C. elegans* and human cell culture suggested that SMG5 induces the UPF1 dephosphorylation via recruitment of the PP2A phosphatase complex^53,60,80,103^. Intriguingly, we find that deleting the C-terminal PIN domain, which was reported to be involved in dephosphorylation of UPF1, completely abolishes the rescue activity of SMG5. It remains to be determined whether the SMG5 PIN domain directly recruits the PP2A complex, as no such interaction was reported in recent SMG5 mass spectrometry experiments^65^. In support of the functional relevance of SMG5 in NMD, it was recently found that the SMG5 homolog in *Drosophila melanogaster* is critical for NMD activity^104^. Of note, no SMG7 homolog exists in *Drosophila*^81,105^, which could explain the strong dependence on SMG5.

The last unclear aspect is how SMG5 gains access to p-UPF1. In normal conditions, this interaction is bridged via SMG7^62^. In the absence of SMG7, a transient interaction between SMG5 and p-UPF1 is likely required not only to activate UPF1 for SMG6-mediated endonucleolytic cleavage, but also to trigger subsequent UPF1 dephosphorylation. In line with this hypothesis, the rescue efficiency of SMG5 14-3-3^mut^ was clearly reduced.

In conclusion, we present here a detailed model for the activation of NMD, which aimed to integrate our insights into the inter-dependency of SMG5, SMG6 and SMG7 with the extensive existing knowledge. This model is centred on UPF1 as molecular mRNA inspector and involves progressive SMG1-mediated phosphorylation as first, and SMG5-SMG7-mediated activation/recycling of UPF1 as second authentication step to identify and degrade NMD targets in a complex transcriptome. The new arrangement of NMD factors in our model creates ample opportunities to investigate their function and interplay and allows the field to move away from earlier models, which were based on parallel or redundant degradation pathways during NMD.

## Online Methods

### Cell lines

Flp-In-T-REx-293 (human, female, embryonic kidney, epithelial; Thermo Fisher Scientific) cells were cultured in high-glucose, GlutaMAX DMEM (Gibco) supplemented with 9% fetal bovine serum (Gibco) and 1x Penicillin Streptomycin (Gibco). The cells were cultivated at 37°C and 5% CO2 in a humidified incubator. The generation of knockout and stable cell lines is described below and all cell lines are summarized in Supplementary Table 4.

### Generation of knockout cells using CRISPR-Cas9

The knockouts were performed using the Alt-R CRISPR-Cas9 system (Integrated DNA Technologies) and reverse transfection of a Cas9:guideRNA ribonucleoprotein complex using Lipofectamine RNAiMAX (Thermo Fisher Scientific) according to the manufacturer’s protocol. The crRNA sequence (Design ID: Hs.Cas9.SMG7.1.AD; Integrated DNA Technologies) to target SMG7 was /AlTR1/rGrArArArArUrGrCrUrArGrUrUrArCrCrGrArUrUrGrUrUrUrUrArGrArGrCrUrArUrGrCrU/AlTR2/. Reverse transfection was performed on 1.5×10^5^ cells per crRNA in 12-well plates. 48 h after transfection the cells were trypsinised, counted and seeded at a mean density of a single cell per well in 96-well plates. Cell colonies originating from a single clone were then screened via Western blot and genome editing of SMG7 was analysed on the genomic level via DNA extraction and Sanger sequencing. Alterations on the transcript level were analysed via RNA extraction followed by reverse transcription and Sanger sequencing.

### DNA and RNA extraction

One day prior to DNA extraction, 2.5×10^5^ cells were seeded in a 6-well plate. To extract DNA, QuickExtract DNA Extraction Solution (Lucigen) was used following the manufacturer’s instructions. RNA was isolated using peqGOLD TriFast (VWR Peqlab) following the manufacturer’s instructions. Following changes were made: Instead of 200 μl chloroform, 150 μl 1-Bromo-3-chloropropane (Molecular Research Center, Inc.) was used. RNA was resuspended in 20 μl RNase-free water.

### Immunoblot analysis

SDS-polyacrylamide gel electrophoresis and immunoblot analysis were performed using protein samples harvested with RIPA buffer (50 mM Tris/HCl pH 8.0, 0.1% SDS, 150 mM NaCl, 1% IGEPAL, 0.5% deoxycholate) or samples eluted from Anti-FLAG M2 magnetic beads. For protein quantification, the Pierce Detergent Compatible Bradford Assay Reagent (Thermo Fisher Scientific) was used. All antibodies (see Key Resources List) were used at the indicated dilutions in 50 mM Tris [pH 7.2], 150 mM NaCl with 0.2% Tween-20 and 5% skim milk powder. Amersham ECL Prime Western Blotting Detection Reagent (GE Healthcare) in combination with the Fusion FX-6 Edge system (Vilber Lourmat) was used for visualization. All antibodies used in this study are listed in Supplementary Table 4. Protein bands detected with the Fusion FX-6 Edge system (Vilber Lourmat) were quantified in a semi-automated manner using the ImageQuant TL 1D software with a rolling-ball background correction. The control condition was set to unity, quantification results are shown as data points and mean.

### Growth assay

To measure growth and mortality rate of cells, CytoTox-Glo Cytotoxicity Assay (Promega) was performed. 10,000 cell/well were seeded in a 96-well plate and assay was performed after 0-5 days using luminometer centro XS3 LB 960 (Berthold Technologies) following the manufacturer’s instructions.

### Stable cell lines and plasmids

The point and deletion mutants of SMG7 were PCR amplified using Q5 polymerase (New England Biolabs) and inserted with an N-terminal FLAG-tag via NheI and NotI (both New England Biolabs) restriction sites into PB-CuO-MCS-IRES-GFP-EF1α-CymR-Puro (System Biosciences). Accordingly, N-terminally FLAG-tagged GST, UPF1 WT and UPF1 SQ N/C-terminal mutant (generated with Integrated DNA Technologies gBlocks and PCR amplification) were cloned via NheI and NotI into PB-CuO-MCS-BGH-EF1-CymR-Puro, which was modified from the original vector by replacing the IRES-GFP cassette with a BGH polyA signal.

The point and deletion mutants of SMG5 were PCR amplified using Q5 polymerase and inserted with an N-terminal FLAG-tag via NheI and NotI restriction sites into the tetracycline inducible pcDNA5/FRT/TO vector (Thermo Fisher Scientific). The mRNA reporter constructs TPI-WT and TPI-PTC160 in the pcDNA5/FRT/TO vector are available on Addgene (IDs 108377-108378).

The cells were stably transfected using the PiggyBac Transposon system (SMG7, UPF1, GST) or using the Flp-In T-REx system (SMG5, mRNA reporter). 2.5-3×10^5^ cells were seeded 24 h before transfection in 6-wells. For PiggyBac stable cells, 2 µg of PiggyBac construct was transfected together with 0.8 µg of the Super PiggyBac Transposase expression vector (System Biosciences) and for Flp-In T-REx stable cells, 1-2 µg of pcDNA5 construct was transfected together with 1 µg of the Flp recombinase expressing plasmid pOG44, using the calcium phosphate method. 48 h after transfection, the cells were transferred into 10 cm dishes and selected with 2 µg ml^−1^ puromycin (InvivoGen) for PiggyBac or 100 µg ml^−1^ hygromycin (InvivoGen) for Flp-In T-REx. After 7-10 days, the colonies were pooled. Expression of the PiggyBac constructs was induced with 30 µg ml^−1^ cumate, Flp-In T-REx constructs were induced with 1 µg ml^−1^ doxycycline. All vectors used in this study are listed in Supplementary Table 4.

Mycoplasma contamination was tested by PCR amplification of mycoplasma-specific genomic DNA^106^ or by using the Mycoplasmacheck service (Eurofins Genomics).

### Reverse transcription, end-point and quantitative RT-PCR

1-4 µg of total RNA was reverse-transcribed in a 20 µl reaction volume with 10 µM VNN-(dT)_20_ primer using the GoScript Reverse Transcriptase (Promega). 2 % of cDNA was used as template in end-point PCRs using the GoTaq Green Master Mix (Promega) or MyTaq Red Mix (Bioline) and 0.2 – 0.6 µM final concentration of sense and antisense primer (see Supplementary Table 4 for sequences). After 30 PCR cycles, the samples were resolved by electrophoresis on ethidium bromide-stained, 1-2% agarose TBE gels and visualized by trans-UV illumination using the Gel Doc XR+ (Bio-Rad). Representative gel images from at least three independent experiments are shown.

Bands detected in agarose gels from the indicated biological replicates of end-point PCRs were quantified using the Image Lab 6.0.1 software (Bio-Rad). Results of the indicated band % per lane are shown as data points and mean. Sanger sequencing of individual bands was performed using the service of Eurofins Genomics.

Quantitative RT-PCR were performed with the GoTaq qPCR Master Mix (Promega) using 2 % of cDNA in 10 µl reactions, 0.2-0.6 µM final concentration of sense and antisense primer (see Supplementary Table 4 for sequences), and the CFX96 Touch Real-Time PCR Detection System (Bio-Rad). The reactions for each biological replicate were performed in duplicates or triplicates and the average Ct (threshold cycle) value was measured. For alternative splicing events, values for canonical isoforms were subtracted from values for NMD sensitive isoforms to calculate the ΔCt. For differentially expressed targets, the values for the housekeeping genes C1orf43 were subtracted from values for the target to calculate the ΔCt. The mean log2 fold changes were calculated from three biologically independent experiments. Log2 fold change results are shown as data points and mean.

### siRNA-mediated knockdowns

Cells were seeded in 6-well plates at a density of 2- 3×10^5^ cells per well and reverse transfected using 2.5 µl Lipofectamine RNAiMAX and 60 pmol of the respective siRNA(s) according to the manufacturer’s instructions. In preparation for 3′ fragment linker ligation, UPF1 phosphorylation and RNA immunoprecipitation (RIP) assays, 3×10^6^ cells were reverse transfected in 10 cm dishes using 6.25 µl Lipofectamine RNAiMAX and 150-200 pmol siRNA. All siRNAs used in this study are listed in Supplementary Table 4.

### RNA-Sequencing and computational analyses

RNA-Seq experiments were carried out with Flp-In-T-REx-293 wild type (WT) cells transfected with Luciferase or SMG5 siRNA and the SMG7 KO clones 2 and 34 transfected with either Luciferase, SMG5 or SMG6 siRNAs. Three biological replicates were analysed for each sample. Total RNA was extracted using peqGOLD TriFast (VWR Peqlab) as described above.

The Lexogen SIRV Set1 Spike-In Control Mix (SKU: 025.03) that provides a set of external RNA controls was added to the total RNA to enable performance assessment. Mix E0 was added to samples with Luciferase siRNA, mix E1 was added to samples with SMG5 siRNA and mix E2 samples with SMG6 siRNA. The Spike-Ins were used for quality control purposes, but not used for the final analysis of DGE, DTU or AS.

The library preparation was performed with the TruSeq mRNA Stranded kit (Illumina). After poly-A selection (using poly-T oligo-attached magnetic beads), mRNA was purified and fragmented using divalent cations under elevated temperature. The RNA fragments underwent reverse transcription using random primers. This is followed by second strand cDNA synthesis with DNA Polymerase I and RNase H. After end repair and A-tailing, indexing adapters were ligated. The products were then purified and amplified to create the final cDNA libraries. After library validation and quantification (Agilent tape station), equimolar amounts of library were pooled. The pool was quantified by using the Peqlab KAPA Library Quantification Kit and the Applied Biosystems 7900HT Sequence Detection System and sequenced on an Illumina NovaSeq6000 sequencing instrument and a PE100 protocol.

Reads were aligned against the human genome (version 38, GENCODE release 33 transcript annotations supplemented with SIRVomeERCCome annotations from Lexogen; obtained from https://www.lexogen.com/sirvs/download/) using the STAR read aligner (version 2.7.3a)^107^. Transcript abundance estimates were computed with Salmon (version 1.1.0)^108^ with a decoy-aware transcriptome. After the import of transcript abundances, differential gene expression analysis was performed with the DESeq2^109^ R package with the significance thresholds |log2FoldChange|> 1 and adjusted p-value (padj) < 0.05. Differential splicing was detected with LeafCutter (version 0.2.8)^110^ with the significance thresholds |deltapsi| > 0.1 and adjusted p-value (p.adjust) < 0.05. Differential transcript usage was computed with IsoformSwitchAnalyzeR (version 1.6.0) and the DEXSeq method^75,111–115^. Significance thresholds were |dIF| > 0.1 and adjusted p-value (isoform_switch_q_value) < 0.05.

PTC status of transcript isoforms with annotated open reading frame was determined by IsoformSwitchAnalyzeR using the 50 nt rule of NMD^75,116–118^. Isoforms with no annotated open reading frame in GENCODE were designated “NA” in the PTC analysis.

The control, SMG7 and SMG6/7 knockdown datasets (Gene Expression Omnibus, GEO: GSE86148)^14^ and UPF1 RIP-seq datasets (GEO: GSE69586)^48^ were processed and analysed with the same programs, program versions, and scripts as the SMG7 KO dataset, with minor changes due to the different sequencing method (paired-end vs. single-end) or lower replicate numbers. For open reading frame prediction, QTI-seq data (NCBI Sequence Read Archive accession SRA160745)^119^ were processed with the Ribo-TISH toolkit^120^. All major parameters for the RNA-Seq analysis are listed in Supplementary Table 4. Overlaps of data sets were represented via nVenn^121^ or the ComplexHeatmap package^122^. Barcode plots were produced with the barcodeplot function from the limma package^113^ using transcript isoform dIF as ranking statistic. Integrative Genomics Viewer (IGV)^123^ snapshots were generated from mapped reads (BAM files) converted to binary tiled data (tdf), using Alfred^124^ with resolution set to 1 and IGVtools. Mean junction counts were obtained from sashimi plots generated using ggsashimi^125^.

### Protein structure modelling and visualization

The structure of human SMG7 (PDB: 1YA0) was superimposed onto the *C. elegans* SMG7 (PDB: 3ZHE) using the MatchMaker command in Chimera version 1.13^126^, to generate a hsSMG7-ceSMG5 hybrid model. ChimeraX version 1.0^127^ was used to visualize the modelled structure.

### Northern blotting

The cells were harvested in peqGOLD TriFast reagent (VWR) and total RNA extraction was performed as described above. 2-4 μg of total RNA were resolved on a 1% agarose/0.4 M formaldehyde gel using the tricine/triethanolamine buffer system^128^ followed by a transfer on a nylon membrane (Roth) in 10x SSC. The blots were incubated overnight at 65°C in Church buffer containing α-32P-GTP [800 Ci/mmol, 10 mCi/ml] body-labelled RNA probes for detection of the reporter mRNA^77^.

Endogenous 7SL RNA was detected by a 5′-32P-labeled oligonucleotide (5′- TGCTCCGTTTCCGACCTGGGCCGGTTCACCCCTCCTT-3′) for which γ-32P-ATP [800 Ci/mmol, 10 mCi/ml] was used for labelling. For NOP56 northern blots, the ex8b riboprobe sequence (5′-GAAACUUGGUCCCUUUGCUGGGCCCUGGGAAUCACUCAGACACCAGGACUGGCCAUCACCCCCAUAGCAGAGGCCUGUAUAGGUCAGGGAGCCCUGGUCAGCCAUCACCGUGAUCCCCCAACAAGCAGUGGGCACCAGAAGUGGCACCUGAUU -3′)^78^ was cloned into the pSP73 vector, linearized and in vitro transcribed using α-32P-GTP [3000 Ci/mmol, 10 mCi/ml]. Ethidium bromide stained 28S and 18S rRNA served as loading controls. RNA signal detected with the Typhoon FLA 7000 (GE Healthcare) was quantified in a semi-automated manner using the ImageQuant TL 1D software with a rolling-ball background correction. EtBr-stained rRNA bands were quantified with the Image Lab 6.0.1 software (Bio-Rad). Signal intensities were normalized to the internal control (7SL or rRNA) before calculation of mean values. The control condition was set to unity (TPI WT for reporter assays), quantification results are shown as data points and mean.

### RNA immunoprecipitation (RIP)

The RIP protocol was adapted from^48^. After seeding and reverse transfecting cells with siRNA, the expression of FLAG-tagged proteins was induced 24 h later. After another 48 h, the cells were washed with 2 ml PBS, harvested in 1 ml PBS, collected for 10 min at 100 xg and resuspended in 300 µl RIP lysis buffer (10 mM Tris pH 7.5, 150 mM NaCl, 2 mM EDTA, 0.1% Triton X-100) supplemented with 1 tablet of PhosSTOP (Roche), 100 µl EDTA-free HALT Protease & Phosphatase Inhibitor (ThermoFisher) and 20 µl RNasin (Promega) per 10 ml buffer. Protein concentration was measured, adjusted to 1 mg total protein in 333 µl RIP buffer, 33 µl of sample was added to 500 µl peqGOLD TriFast and saved as input sample. The remaining sample was combined with 35 µl pre-washed Anti-FLAG M2 Magnetic Beads (Sigma-Aldrich). The samples were incubated in an overhead rotator for 2 h at 4 °C, washed 5x with 1 ml RIP Wash Buffer (5 mM Tris pH 7.5, 150 mM NaCl, 0.1% Triton X-100) and co-immunoprecipitated RNA was recovered by incubating the beads with 500 µl peqGOLD TriFast for 10 min at room temperature.

Both input and IP samples were subjected to RNA extraction with the addition of 1 µl Precipitation Carrier. 10 µl of resuspended RNA was used for reverse transcription and 2 % of cDNA was used for quantitative PCR.

### Co-immunoprecipitation

FLAG-tagged proteins were expressed in stable cell lines (2.5-3.0 x 10^6^ cells per 10 cm dish) induced for 48-72 h. When indicated, the cells were subjected to an okadaic acid (OA) treatment for 2 h before harvesting with 50 or 100 nM OA. The samples were lysed in buffer E (20 mM HEPES-KOH (pH 7.9), 100 mM KCl, 10% glycerol, 1 mM DTT, Protease Inhibitor, 1 µg ml^−1^ RNase A) and sonicated using the Bandelin Sonopuls mini20 with 15 pulses (2.5 mm tip in 600 µl volume, 1s pulse, 50% amplitude). Concentration-adjusted lysates were subjected to immunoprecipitation for 2 h using Anti-FLAG M2 Magnetic Beads (Sigma-Aldrich), the beads were washed three times for 5 min with buffer E, mild wash buffer (20 mM HEPES-KOH (pH 7.9), 137 mM NaCl, 2 mM MgCl_2_, 0.2% Triton X-100, 0.1% NP-40) or medium wash buffer (20 mM HEPES-KOH (pH 7.9), 200 mM NaCl, 2 mM MgCl_2_, 0.2% Triton X-100, 0.1% NP-40, 0.05% Na-deoxycholate). Co-immunoprecipitated proteins were eluted with SDS-sample buffer, separated by SDS-PAGE, and analysed by immunoblotting.

### Data Availability

The accession number for the raw RNA-sequencing data reported in this paper is ArrayExpress: E-MTAB-9330. The authors declare that all the data supporting the findings of this study are available within the article and its Supplementary Information files and from the corresponding authors upon reasonable request.

### Code Availability

For availability of codes that were developed for this project, please contact the corresponding authors.

## Acknowledgements

We thank members of the Gehring lab for discussions and reading of the manuscript. We also thank Marek Franitza and Christian Becker (Cologne Center for Genomics, CCG) for preparing the sequencing libraries and operating the sequencer. We acknowledge Tobias Jakobi for helping with infrastructure support. This work was supported by grants from the Deutsche Forschungsgemeinschaft to C.D. (DI 1501/8-1, DI1501/8-2) and N.H.G (GE 2014/6-2 and GE 2014/10-1). V.B. was funded under the Institutional Strategy of the University of Cologne within the German Excellence Initiative. N.H.G. acknowledges support by a Heisenberg professorship (GE 2014/7-1 and GE 2014/13-1) from the Deutsche Forschungsgemeinschaft. C.D. and T.B.B. were kindly supported by the Klaus Tschira Stiftung gGmbH (00.219.2013). This work was supported by the DFG Research Infrastructure as part of the Next Generation Sequencing Competence Network (project 423957469). NGS analyses were carried out at the production site WGGC Cologne.

## Author Contributions

*Conceptualization*, N.H.G., S.K. and V.B.; *Methodology*, V.B., S.K., N.H.G.; *Software*, V.B., T.B.B., J.V.G., and C.D.; *Investigation*, S.K., V.B. and J.V.G.; *Resources and Data Curation*, V.B., T.B.B., J.A., and C.D.; *Writing – Original Draft, Review & Editing*, V.B., J.V.G., N.H.G. and S.K.; *Visualization*, V.B. and S.K.; *Supervision*, N.H.G. and V.B.; *Funding Acquisition*, N.H.G. and C.D.

---

**Extended Data Fig. 1:**
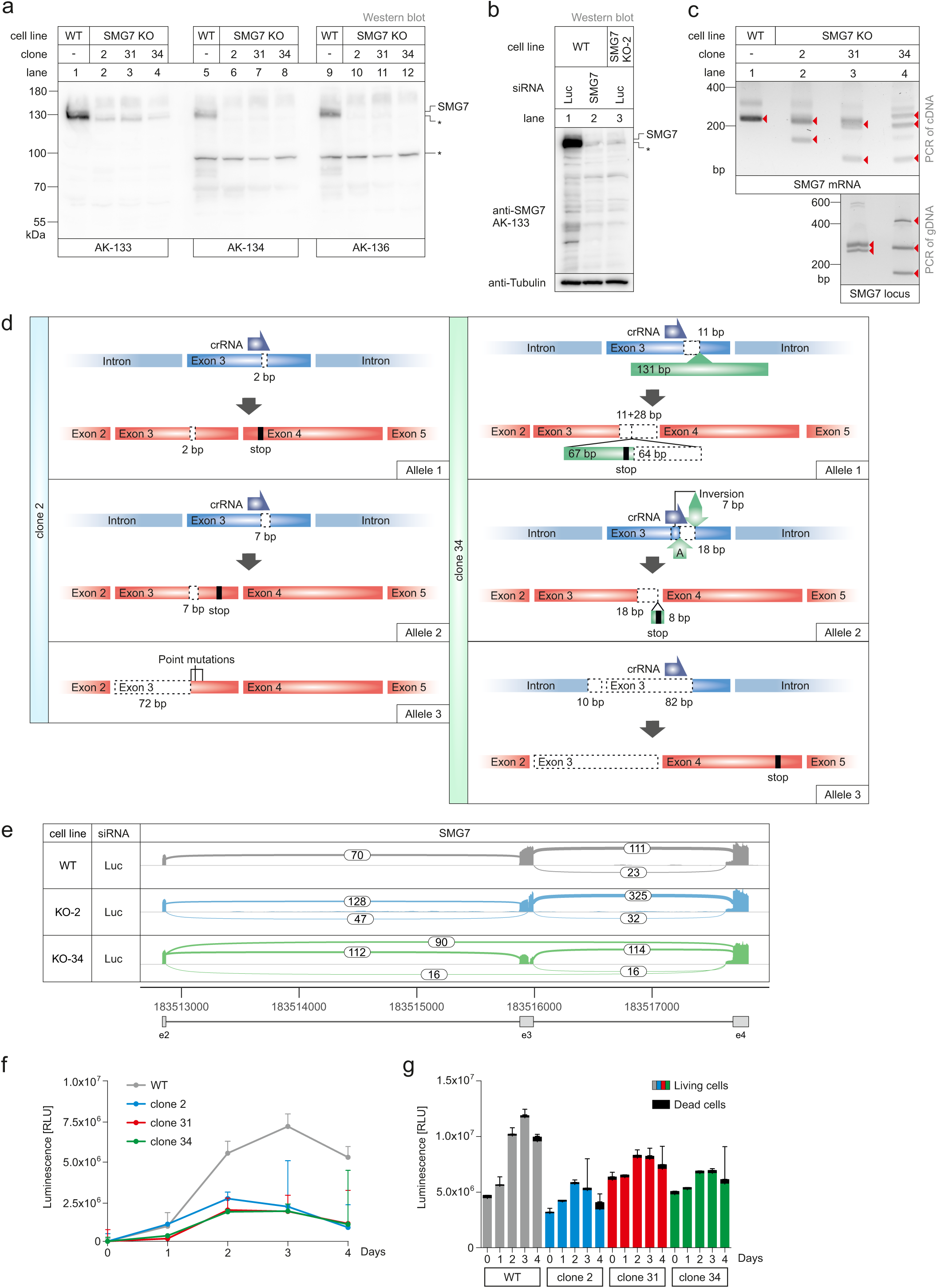
Characterization of SMG7 knockout cells. **a**, Uncropped western blot analysis of SMG7 knockout (KO) cell lines. **b**, Comparison between SMG7 knockout (KO) and knockdown (KD) conditions with AK-133. **c**, Detection of SMG7 genomic alterations and splicing variants via PCR. Red arrows indicate identified bands. **d**, Overview of CRISPR-Cas9 induced alterations on the genomic and transcriptomic level concerning the SMG7 locus. **e**, Sashimi plot of SMG7 WT, KO-2 and KO-34 clones from RNA-seq data, highlighting alternative splicing events. **f**, Luminescence-based growth assay of the indicated cell lines. **g**, Quantification of live and dead cells at the indicated time points of the growth assay.

**Extended Data Fig. 2:**
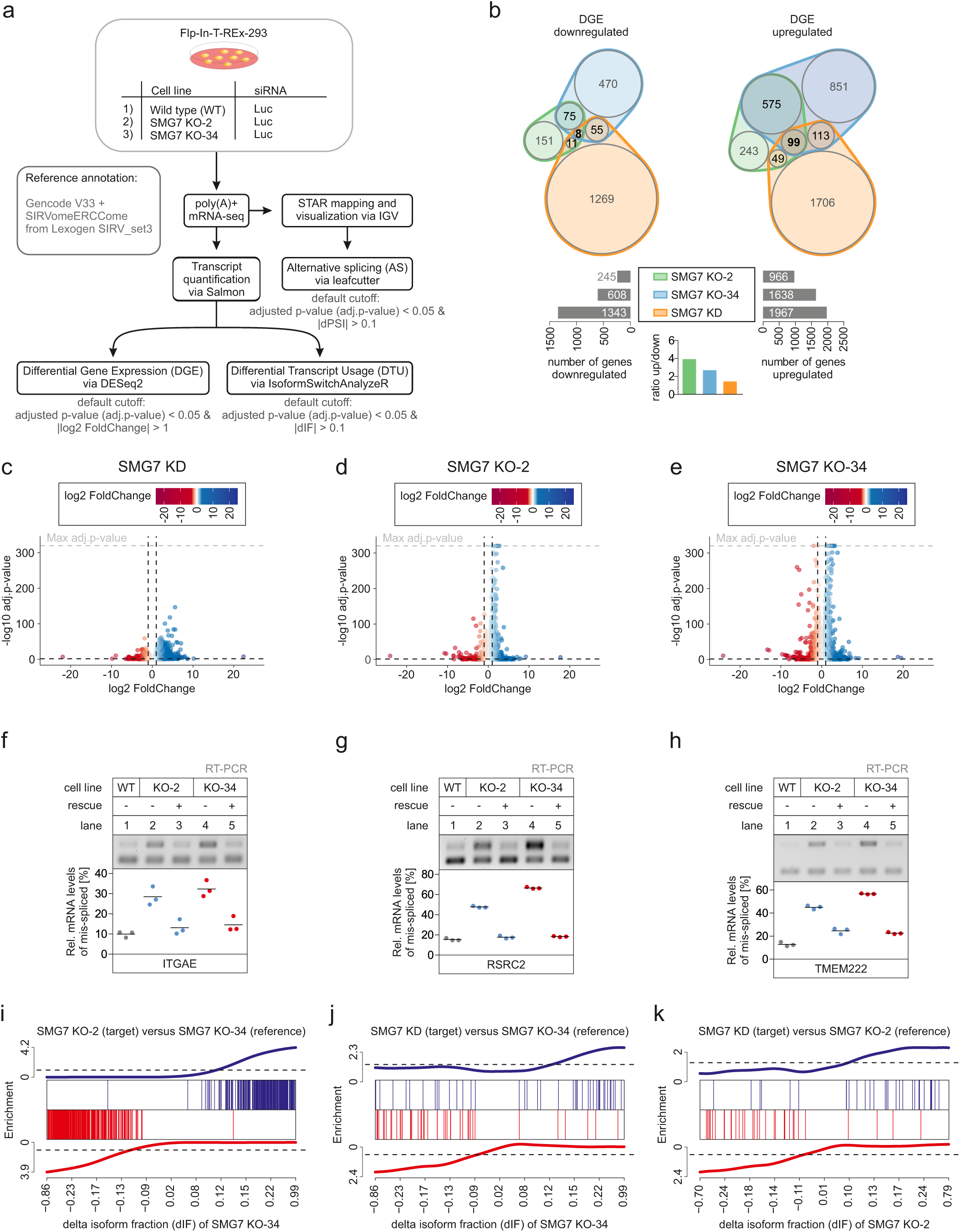
RNA-Seq analyses of SMG7 KO cells. **a**, Schematic overview of the RNA-Seq analysis pipeline. **b**, Overlap of upregulated and downregulated genes in the differential gene expression (DGE) analysis. Total numbers and ratios are given below. **c-e**, Volcano plots showing the differential gene expression analyses from the indicated RNA-seq dataset. **f-h**, End-point RT-PCR detection of ITGAE (F), RSRC2 (G) and TMEM222 (H) transcript isoforms was carried out in the indicated cell lines, with or without expression of FLAG-tagged SMG7 rescue constructs. Data points and means from the gel quantification are plotted (n=3). **i-k**, Barcode plots showing the enrichment of transcript isoforms that undergo isoform switching (adjusted p-value < 0.05) of the target condition compared to the reference dataset indicated on the x-axis. On the x-axis the dIF of transcripts that undergo isoform switching in the reference are ranked according to their dIF.

**Extended Data Fig. 3:**
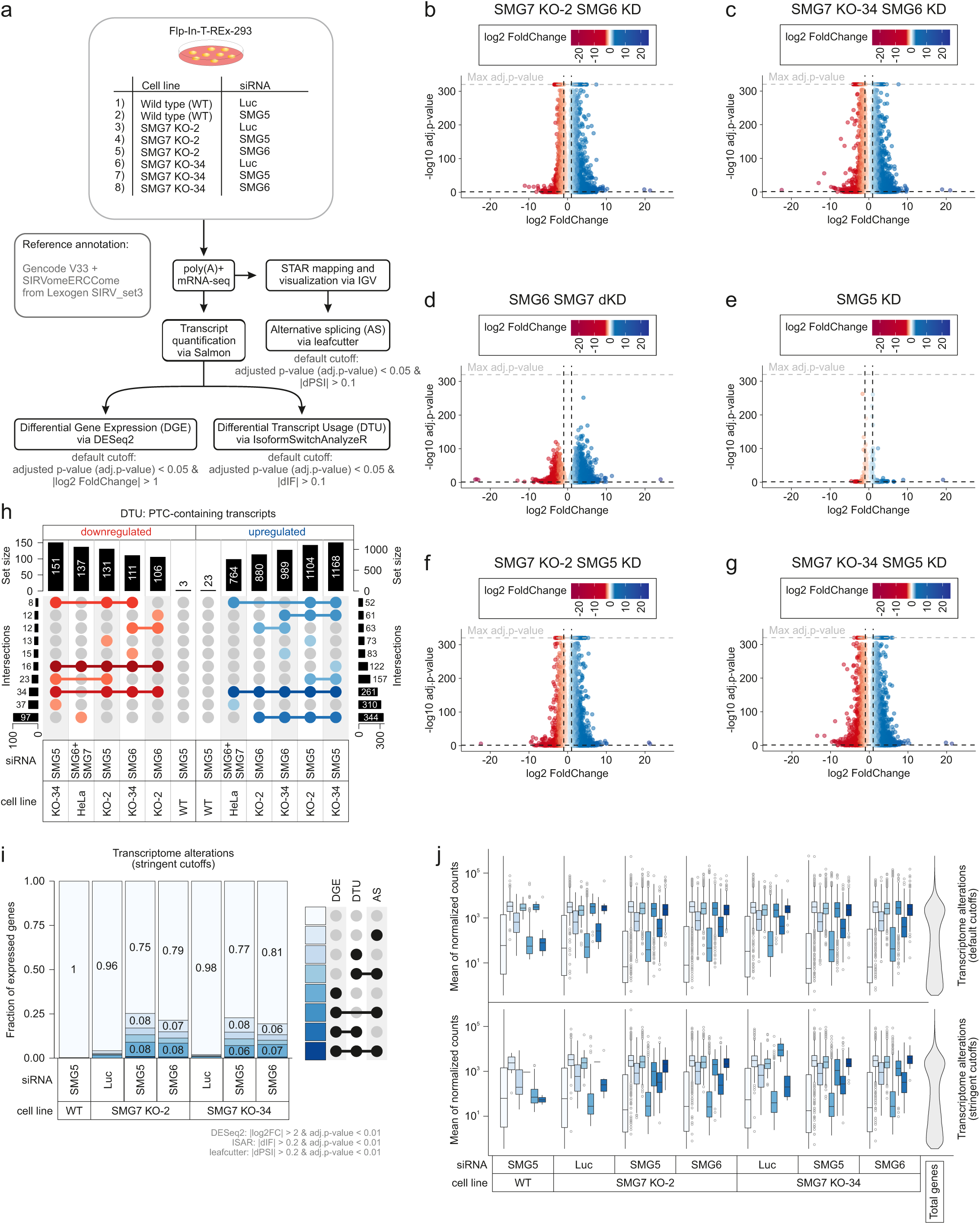
Analyses of SMG7 KO plus knockdown RNA-seq data. **a**, Schematic overview of the analysis pipeline. **b-g**, Volcano plots showing the differential gene expression analyses from the indicated RNA-seq dataset. **h**, Overlap of up- or downregulated premature termination codon (PTC)-containing isoforms in the RNA-seq data is shown as UpSet plot. Only the top 10 overlapping sets are shown. **i**, Fraction of expressed genes (genes with non-zero counts in DESeq2) which undergo differential gene expression (DGE), differential transcript usage (DTU) and/or alternative splicing (AS) with stringent cutoffs. **j**, Boxplots showing the distribution of different combinations of transcriptomic events in relation to the expression of the gene (indicated by the mean of the normalized counts), with default (top) or stringent (bottom) cutoffs. The total distribution of gene expression is shown on the right as violin plot.

**Extended Data Fig. 4:**
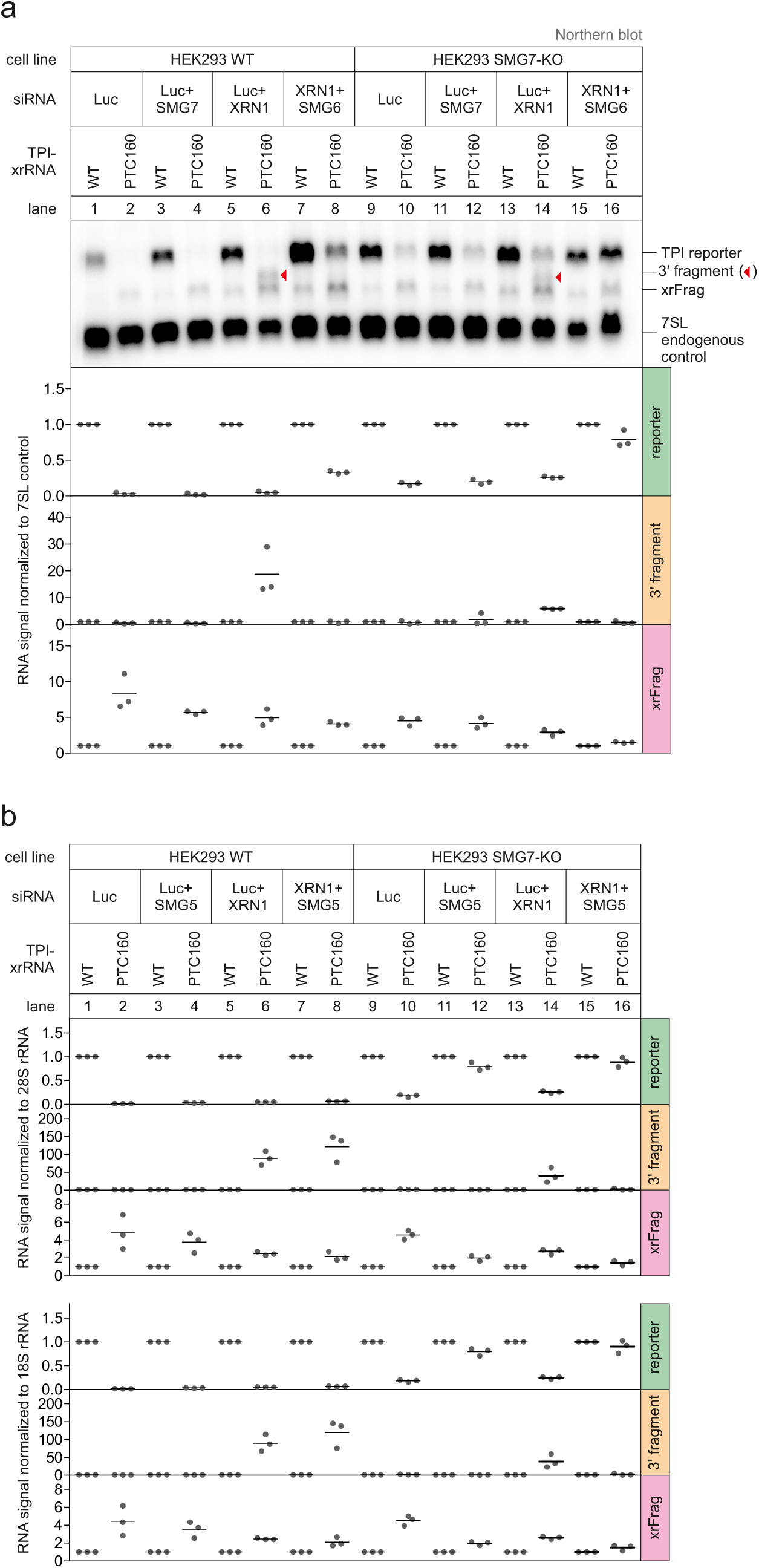
Northern blot analysis of endonucleolytic cleavage. **a**, Northern blot analysis of triosephosphate isomerase (TPI) reporter, 3′ fragments, xrFrags and 7SL endogenous control. Quantification results are shown as data points and mean (n=3). **b**, Quantification of Fig. 4b with 28S or 18S rRNA as reference RNA. Quantification results are shown as data points and mean (n=3).

**Extended Data Fig. 5:**
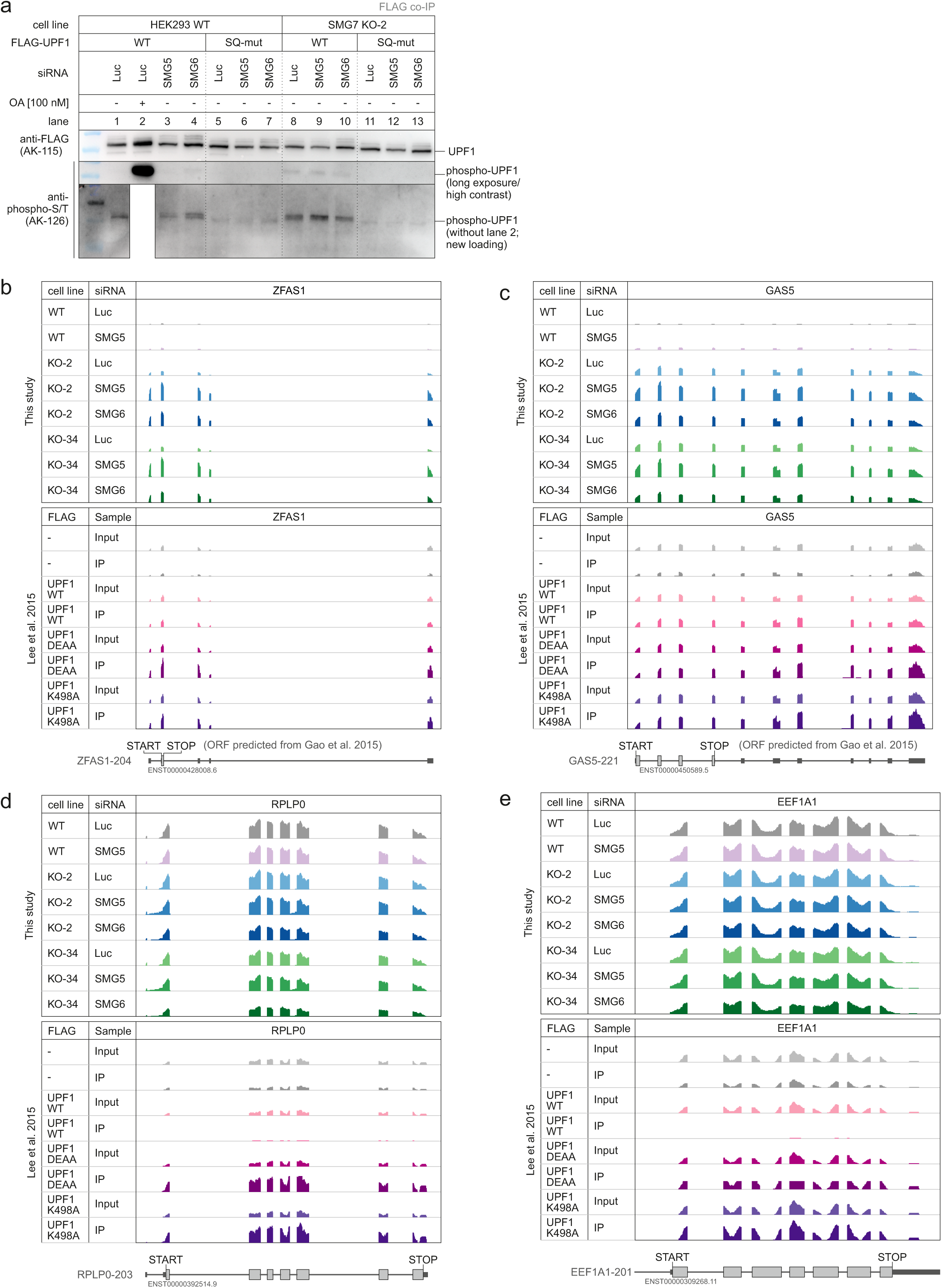
UPF1 hyper-phosphorylation and target discrimination. **a**, Analysis of UPF1 phosphorylation status after co-immunoprecipitation (IP) of expressed FLAG-tagged UPF1 WT or SQ mutant. Okadaic acid (OA) served as inhibitor of PP2A to prevent UPF1 dephosphorylation. **b-e**, Read coverage of the indicated genes from SMG7 KO plus knockdown RNA-seq data and published RIP-Seq data (Lee et al. 2015) are shown as Integrative Genomics Viewer (IGV) snapshots. The canonical isoform is schematically indicated below. If no translated ORF was annotated, QTI-seq data (Gao et al. 2015) were used to predict ORFs.

